# US-align: Universal Structure Alignments of Proteins, Nucleic Acids, and Macromolecular Complexes

**DOI:** 10.1101/2022.04.18.488565

**Authors:** Chengxin Zhang, Morgan Shine, Anna Marie Pyle, Yang Zhang

## Abstract

Structure comparison and alignment are of fundamental importance in structural biology studies. We developed the first universal platform, US-align, to uniformly align monomer and complex structures of different macromolecules (proteins, RNAs, and DNAs). The pipeline is built on a uniform TM-score objective function coupled with a heuristic alignment searching algorithm. Large-scale benchmarks demonstrated significant advantages of US-align over state-of-the-art methods in pairwise and multiple structure alignments of different molecules. Detailed analyses showed that the major advantage of US-align lies in the extensive optimization of the unified objective function powered by efficient heuristic search iterations, which significantly improve the accuracy and speed of the structural alignment process. Meanwhile, the universal protocol fusing different molecular and structural types helps facilitate the heterogeneous oligomer structure comparison and template-based protein-protein and protein-RNA/DNA docking.

## Introduction

Structural comparison and alignment of biomacromolecules, including protein, RNA, and DNA, are of fundamental importance in structural biology studies. Apart from providing intuitive visualizations of the shape comparisons, structure alignment is needed for structure-based protein function annotation^1-3^, modeling mutation effects^4^, rational protein design^5, 6^, and protein structure classification^7^. Recent applications have also been seen in the use of templates identified by structure alignment for inter-domain structural assembly^8^ and template-based protein-RNA docking^9^.

Different methods have been developed for comparing different types of molecules. For example, Dali^10^ and TM-align^11^ are typical algorithms to align protein monomer structures by maximizing both alignment accuracy and coverage (the portion of aligned residues divided by the sequence length). Similarly, RNA-align^12^, RMalign^13^, STAR3D^14^, and ARTS^15^ were designed for aligning RNA and DNA molecules, while MM-align^16^ was proposed to compare multi-chain protein complex structures. Recently, algorithms such as mTM-align^17^, Matt^18^, and MUSTANG^19^ were proposed for aligning multiple protein structures. Despite their usefulness, choosing an algorithm suitable for a specific molecular alignment task can be confusing for biological users. Meanwhile, the use of different assessment matrices for different methods makes the mutual structural comparisons of different molecule types difficult.

The most widely used structural comparison matrix is the root mean square deviation (RMSD)^20^ of two molecule structures. It is however not suitable for structure alignment because minimizing RMSD of structurally aligned regions often results in low alignment coverage. GDT^21^ and MaxSub^22^ were later proposed to optimize alignment accuracy and coverage simultaneously. However, both GDT and MaxSub scores are sequence length dependent, as the average score for random structure pairs has a power-law dependence on the sequence length^23^, which renders the absolute magnitude of these scores meaningless. To address these issues, TM-score was proposed as the first size-independent metric by the introduction of a length-dependent scale 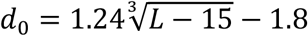 to normalize the residue distance^23, 24^, i.e., 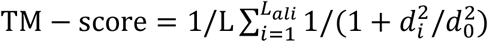, where *L* is the length of the target structure; *L*_*ali*_ is the number of aligned residue pairs; and *d*_*i*_ is the distance between the Cα atoms of the *i*-th pair of aligned residues. The TM-score was recently extended to TM-score_RNA_ for nucleic acid structure comparison^12^ (see **Supplementary Text 1** for a complete discussion on TM-score and TM-score_RNA_). The TM-score was recently extended to TM-score_RNA_ for nucleic acid structure comparison^12^. The unification of scoring function provides the potential to unify the structural comparison of different molecules and molecular complexes.

In this work, we developed a Universal Structure Alignment (US-align) platform, which performs 3D structure alignments for monomeric and complex protein and nucleic acid structures, built on the well-established TM-score and heuristic structural alignment algorithms. The universal strategy to address all macromolecular structure alignments makes the alignments of heterogeneous complexes (such as protein-RNA complexes) feasible. Meanwhile, the extensive optimization of a uniform scoring metric enables the algorithm to generate faster and more accurate alignments compared to the state-of-the-art methods developed for specific structural alignment tasks. The source code and the on-line server of US-align are freely available at https://zhanggroup.org/US-align/, which accepts both legacy PDB and mmCIF/PDBx formats^25^ and automatically recognizes and selects the optimized algorithms for different input structure types.

## Results

US-align is a versatile structural alignment program that performs four different modes of alignments, each of which can handle structures of proteins, RNAs, and DNAs (**Fig. 1**). The first mode is monomeric structure alignment (**Fig. 1a**) by establishing the residue-level correspondences with optimal superimposition between a pair of monomeric chains. The second is oligomeric alignment (**Fig. 1b**), which establishes both chain-level and residue-level correspondences between a pair of oligomeric structures, each with two or more chains. The third is for multiple structure alignment (MSTA, **Fig. 1c**), where the input consists of three or more monomeric structures and the output is a consensus alignment among all structures. The last mode of US-align is template-based docking (**Fig. 1d**), which assembles two or more individual chains together by matching them to an oligomer template. The core idea of US-align is built on the construction of multiple heuristic alignments that cover different initial postures to avoid the trap of a specific local minimum, an issue suffered by many structural alignment methods. The follow-up rapid dynamic programming iterations help improve both accuracy and speed of the alignment procedures. The following sections benchmark the performance of US-align on the four different alignment tasks.

**Fig. 1.**
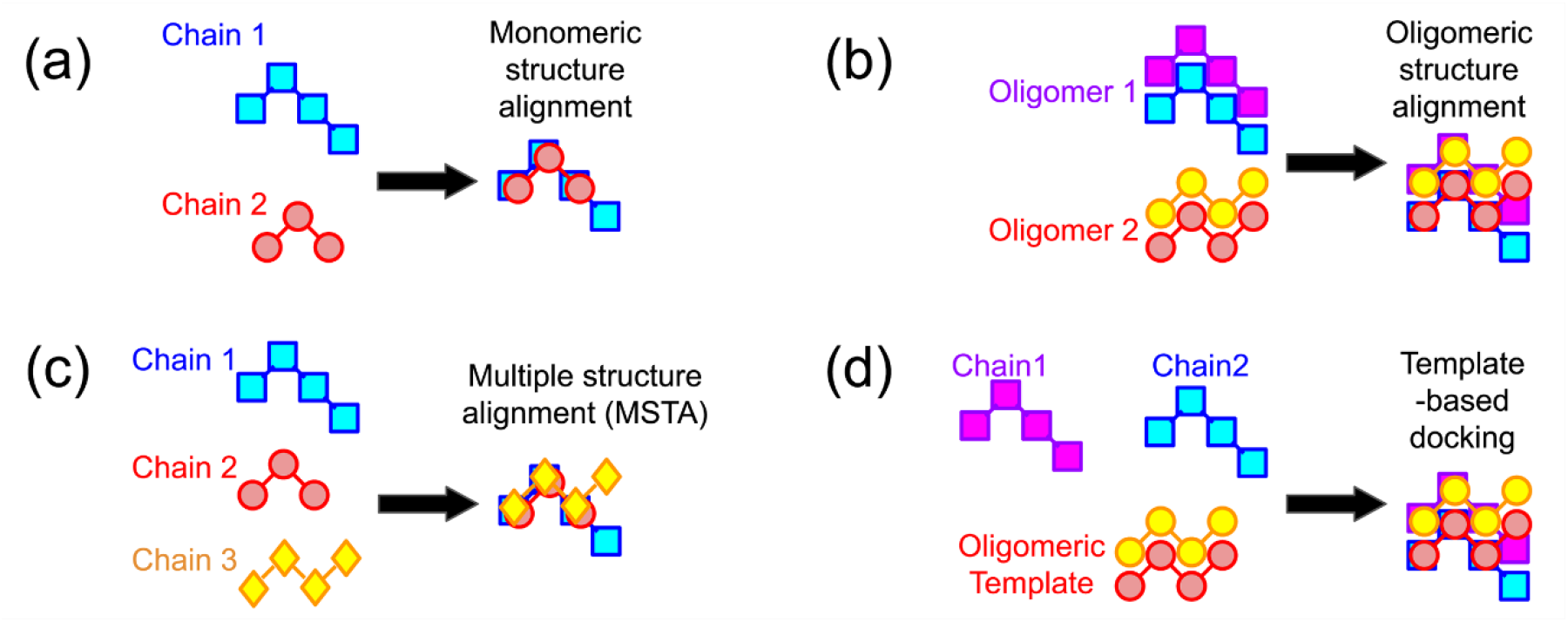
Different structure alignment modes of US-align, which can perform **a**, pairwise monomeric structure alignment; **b**, pairwise oligomeric structure alignment; **c**, MSTA; and **d**, template-based docking of monomeric chains into an oligomeric structure. Different chains are distinguished by different colors and marker styles in this schematic.

### Oligomeric Structure Alignment

We first benchmarked US-align against two open-source programs for oligomeric structure alignments, MM-align^16^ and MICAN^26^ for oligomeric structure alignments. While MM-align generates structure alignments by exhaustive combination of TM-align alignments for each individual chain pair, MICAN is built on a hierarchical strategy of secondary structure element (SSE) and residue-level alignments. The three programs were benchmarked on a set of 1,123 protein complex structures collected from the PDB that are non-redundant at a pairwise sequence identity cutoff of 30%. The dataset includes 200 dimers, 200 trimers, 200 tetramers, 129 pentamers, 200 hexamers, 60 heptamers and 134 octamers (described in detail in Supplementary **Text 2**).

**Fig. 2** summarizes the performance of the three oligomeric alignment programs in terms of TM-score, RMSD, alignment coverage, and execution time for all-against-all alignments among the structures with the same number of chains. As TM-score and coverage for the alignment of the same pair of structures could differ depending on whether the TM-score and coverage were normalized by the longer or the shorter structure, we reported the TM-score and coverage normalized by the shorter structure for the remainder of this manuscript, unless mentioned otherwise.

**Fig. 2.**
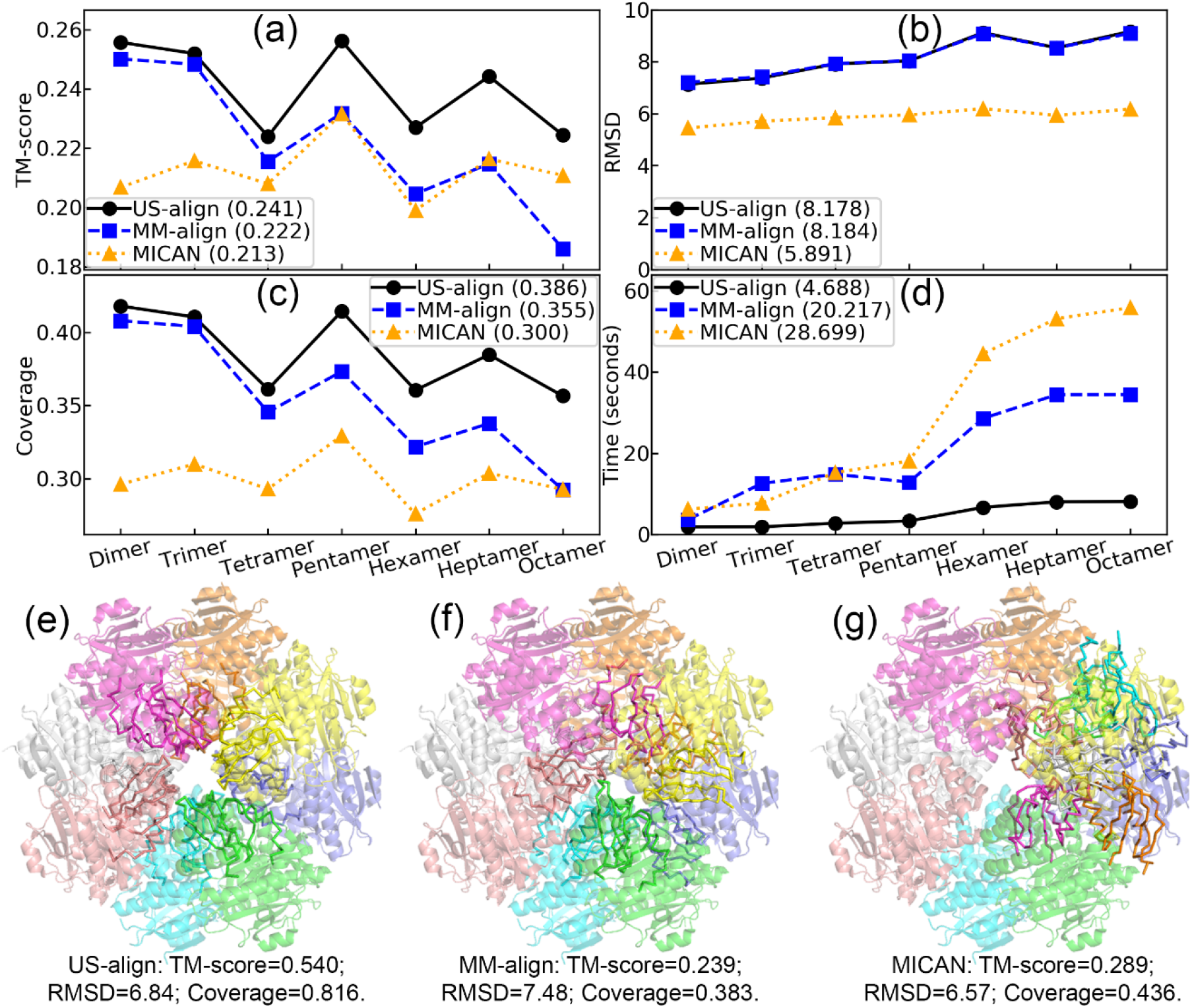
Performance of oligomeric protein structure alignment by US-align, MM-align, and MICAN in terms of average **a**, TM-score, **b**, RMSD, **c**, alignment coverage, and **d**, running time for complex structures in different oligomeric states (*x*-axis). The standard error of mean (SEM) values for all metrics are comparable across different methods and very small **(Supplementary Table 1**). Therefore, the error bars for SEM are invisible. **e-g**, Octamer alignment between PDB 4JHM (semi-transparent cartoon) and PDB 4IAJ (ribbon). Each chain of the oligomer is shown in a different color.

The data show that US-align consistently outperformed both MM-align and MICAN on TM-score, coverage, and execution time. However, it does appear that MICAN has a lower RMSD compared to US-align and MM-align in **Fig. 2b**. This is because MICAN alignment covers a much smaller portion of the full structure than the other methods (**Fig. 2c**), which is also the reason for the low TM-score of MICAN due to the lack of balance between alignment accuracy and coverage. The average TM-score of US-align across all types of oligomers (0.243) is 8.6% and 13.1% higher than those of MM-align (0.224) and MICAN (0.215), which correspond to p-values<1E-303 by Student’s t-test.

It is noted that the performance of a structural alignment method usually relies on both the alignment search engine and the objective function. The difference shown here is apparently not caused by the scoring functions, as US-align, MM-align, and MICAN in this benchmark all use TM-score as the objective function. Therefore, these data highlight the efficiency of the heuristic searching process in US-align which covers larger and more important spaces of chain assignments and structural alignments in a limited amount of CPU time. This difference was particularly evident for oligomers with more chains. For example, the average TM-score of US-align was only 2.2% higher than MM-align for the dimers but 20.6% higher than MM-align for the octamers (**Supplementary Table 1**). One reason for these performance differences for larger oligomers was the better ability of US-align to identify correct chain correspondences, especially for oligomers with high symmetry. If we count all 134 octamers that are the octamers of the highest number of chains in our test dataset, for example, US-align generated alignments containing on average 6.8 aligned chain pairs, which was 25.9% and 13.3% higher than those from MM-align (5.4) and MICAN (6.0), respectively. **Fig. 2e-g** show and example of the octamers from the mandelate racemase/muconate lactonizing enzyme (PDB 4JHM) and the SP_1775 protein (PDB 4IAJ) with D_4_ symmetry. The optimal alignment, derived by US-align with TM-score=0.540, aligned each of the 8 chains in 4JHM to one chain in 4IAJ (**Fig. 2e**). On the other hand, MM-align (**Fig. 2f**) and MICAN (**Fig. 2g**) only aligned 5 and 3 out of the 8 chains, respectively, leading to much lower TM-scores of 0.239 and 0.289, respectively. Although US-align generates on average more accurate oligomeric structural alignments, it could still generate suboptimal chain assignments for 2% of the cases in our test set, where one example is given in **Supplementary Fig. 1** for which US-align underperforms MM-align. This is mainly because the initial chain assignment by US-align is generated by a heuristic search algorithm: the Enhanced Greedy Search (EGS; see Methods section). Although EGS significantly improves the speed of chain assignment with little compromise in accuracy in general, it may still very occasionally miss the best chain assignments that could otherwise be detected by an exhaustive search, as that implemented by MM-align.

As a unique advantage of the universal structure alignment approach, US-align can perform oligomeric alignments for nucleic acid-nucleic acid or protein-nucleic acid complexes, while both MM-align and MICAN could only deal with protein-protein complexes. In **Fig. 3**, we present a case study of structure alignments between two protein-RNA complexes from two different bacteria. Since the protein components of both complexes (PDB 1Y39 chain A and PDB 2ZJR chain F) are 50S ribosomal proteins L11, they share a high structural similarity (TM-score=0.784, **Fig. 3a**). Similarly, the RNA components of the two complexes (PDB 1Y39 chain C and 2ZJR chain X) are fragment and full-length 23S rRNAs, respectively, and share a high similarity (TM-score=0.785, **Fig. 3b**). When combining them together, US-align creates an alignment with an even higher similarity (TM-score=0.861, **Fig. 3c**) due to the cooperative optimization of the complex alignments. In **Supplementary Fig. 2**, we show another example where the hetero-oligomeric alignment by US-align between a protein-DNA complex and a protein-RNA complex revealed a similar mode of interaction with a significant TM-score of 0.467, which could not otherwise be captured by monomeric alignments (TM-score=0.301 and 0.157, respectively, both below the statistical significance threshold).

**Fig. 3.**
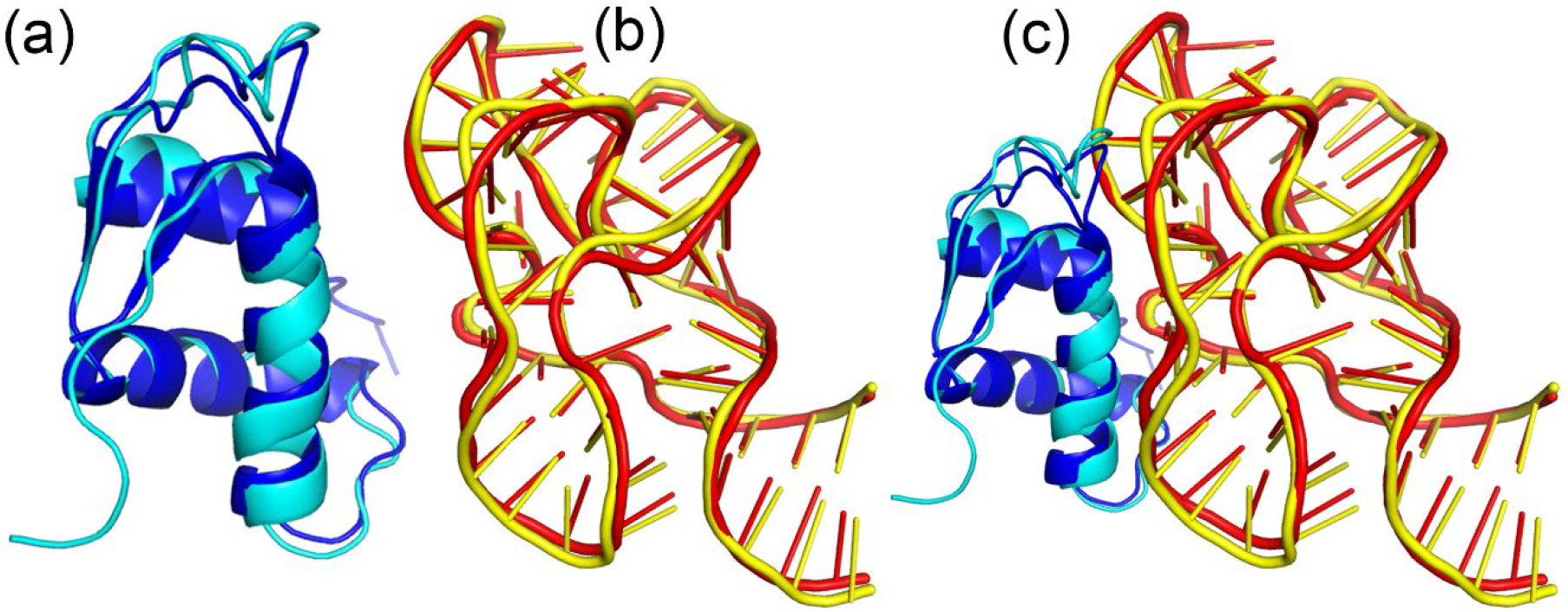
US-align alignment between PDB 1Y39 (chain A in blue; chain C in red) and PDB 2ZJR (chain F in cyan; chain X in yellow). **a**, Alignment of the protein components. **b**, Alignment of the RNAs. **c**, Alignment of the full protein-RNA complexes. Since the full 2ZJR chain X has 2,686 residues and is too large to show in the figure, panels **b** and **c** only show residues 1,063 to 1,119, which correspond to the region aligned to 1Y39 chain C.

### Monomeric Structure Alignment

Structural alignments on single-chain monomer structures are a fundamental component of US-align. To examine the effectiveness of RNA monomer structure comparisons, we first used CD-HIT-EST^27^ to cluster the sequences of all 3,724 unique RNA chains from the PDB, resulting in 637 chains with sequence length ≥ 30 nucleotides and pairwise sequence identity<80%. We then ran an all-against-all pairwise alignment of these 637 chains by US-align, together with four other programs: RMalign^13^, STAR3D^14^, ARTS^15^, and Rclick^28^. The data in **Fig. 4a-d** and **Supplementary Table 2** show that US-align outperforms all four control RNA structure alignment programs with TM-score_RNA_ 5.8% higher than RMalign, 27.5% higher than STAR3D, 34.5% higher than ARTS, and 38.6% higher than Rclick, where the difference corresponds to p-value<1E-303 for all TM-score comparisons. Meanwhile, USalign is 9.6, 31.6, 2.0, and 45.7 times faster than the four control programs, respectively.

**Fig. 4.**
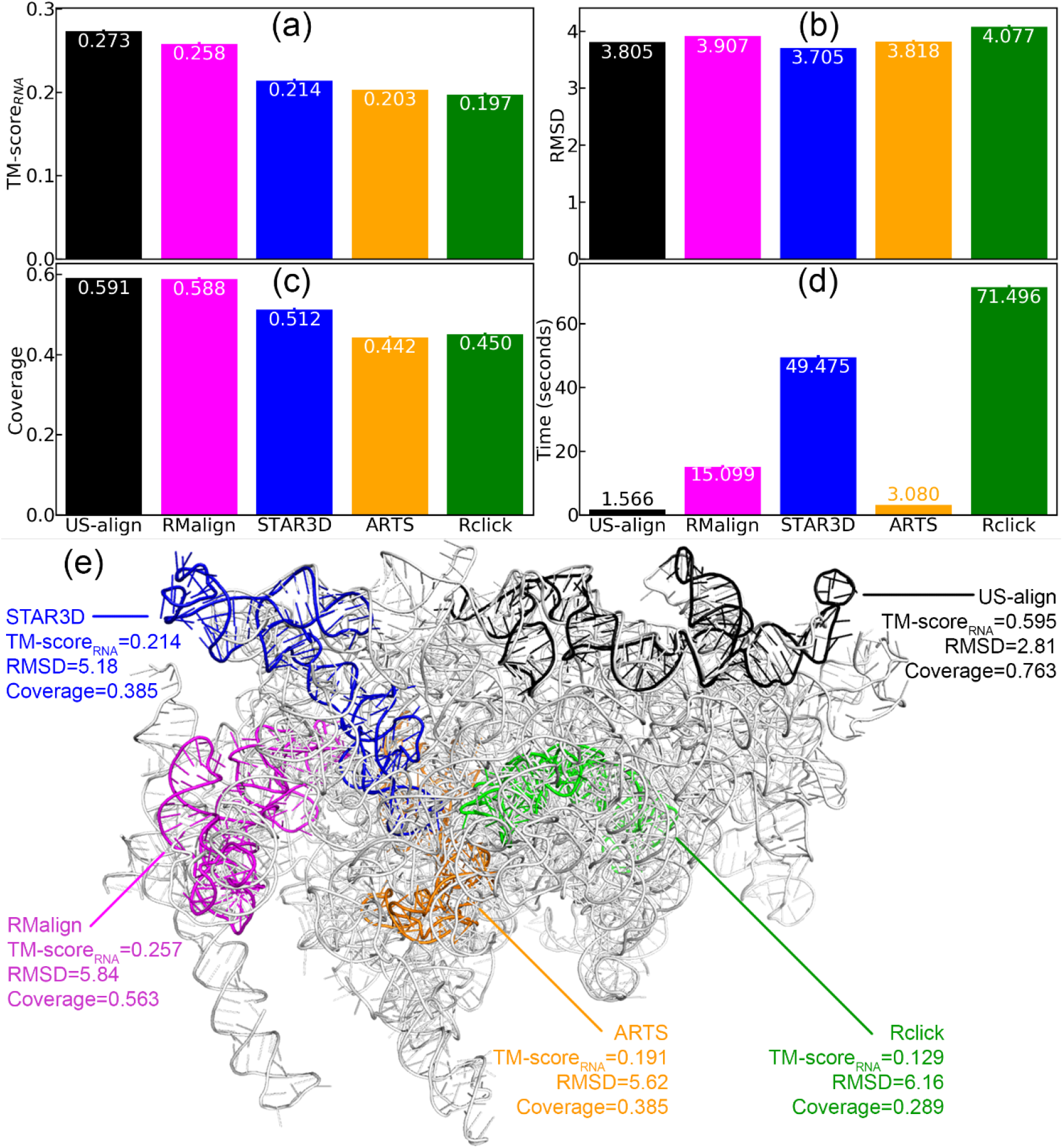
**a-d**, Overall performance of pairwise structure alignment of monomeric RNA chains by US-align and third-party programs, including RMalign, STAR3D, ARTS, and Rclick. Since STAR3D, ARTS, and Rclick only generated results for 86.8%, 84.0%, and 99.8% of all pairs, respectively, this figure is for the subset of 168,917 chain pairs for which all programs have alignment results. The performance was measured by **a**, TM-score_RNA_, **b**, RMSD, **c**, coverage, and **d**, running time. Error bars on top of most bars were not visible because the standard error of mean (SEM) values for all metrics were close to zero (**Supplementary Table 2**). **e**, Monomeric RNA structure alignments generated by different methods between a pair of eukaryotic rRNAs from the large ribosomal subunit: short rRNA-IV (PDB ID 4V8M chain BH, 135 nucleotides; colored based on alignment programs) and 28S rRNA (PDB ID 6Y2L chain L5, 3,613 nucleotides; white), with a low pairwise sequence identity of 20%.

In **Fig. 4e**, we ran the RNA structural alignment programs to match a short rRNA-IV (PDB ID 4V8M with 135 nucleotides) with a large 28S rRNA (PDB ID 6Y2L chain L5, 3,613 nucleotides). Only US-align could identify the correct alignment with a TM-score=0.595, which is 2.3 to 4.6 times higher than that identified by the other four programs. This example highlights the ability of US-align to handle RNA structure pairs with complex topologies and low sequence identities (20% in this example).

In **Supplementary Fig. 3**, we summarized the comparison results of US-align with 4 state-of-the-art monomeric protein structure alignment methods: SPalign^29^, Dali^10^, MICAN^26^, and SSM^30^. On the 31,951 pairs of protein structures, which were collected from the all-to-all pairing of a subset of 1,000 proteins from the SCOPe 2.06, US-align creates alignments with a reasonable combination of RMSD (4.546 Å) and coverage (68.9%), resulting in the highest TM-score (0.447), which is 2.1%, 8.2%, 13.2%, and 21.5% higher than that achieved by SPalign, Dali, MICAN, and SSM, respectively, with a *p*-value ≤ 3.40E-26 in all comparisons (**Supplementary Table 3**). If we count for the number of the alignments with a TM-score≥ 0.5 ^24^, US-align identified 8119 pairs of similar global folds, which is 6.0%, 34.0%, 72.0% and 148.4% higher than that by SPalign (7661), Dali (6050), MICAN (4720), and SSM (3268), respectively (**Supplementary Table 4**). Meanwhile, the CPU time of US-align is 2.7, 6.2, 3.0, and 1.6 times lower than the benchmark programs (**Supplementary Fig. 3d**).

US-align also has good performance when evaluated on the objective functions from other programs, such as Q-score and Dali Z-score which are unique to the SSM^30^ and Dali^10^ programs, respectively (see **Supplementary Text 3**). On the same non-redundant SCOPe dataset, US-align achieves the highest average Q-score of 0.105 (**Supplementary Fig. 3e**), which is 38.1%, 118.7%, 22.1% and 98.1% higher than those by SPalign, Dali, MICAN and SSM, respectively, with p-values ≤ 5.93E-242 for all comparisons (**Supplementary Table 4**). Similarly, US-align achieves the second highest Dali Z-score (0.910), which is lower than MICAN (1.593) but significantly higher (with p-value ≤ 7.48E-55, **Supplementary Table 4**) than SPalign (0.375), Dali (−1.894), and SSM (−3.175) (**Supplementary Fig. 3f**). The reason for the higher Dali Z-score by MICAN is that MICAN tends to generate alignments with lower coverage than US-align (by 19.2%). Since Dali Z-score is more sensitive to local variations than TM-score and Q-score, the sacrifice of alignment coverage for a smaller distance deviation at the aligned region results in a more favorable Dali Z-score by MICAN. Overall, the good performance of US-align on a broad range of scoring metrics reinforces the above observations that US-align has not been over-optimized for its own objective function and that its efficient alignment search engine allows it to derive reasonable alignments in a generic sense.

As an alternative assessment on the alignment performance, we calculated the agreements of manually created pairwise protein alignments from the MALIDUP dataset^31^ and those from automatic protein structure alignment (**Supplementary Fig. 4**, and **Supplementary Table 5**). Here, to avoid overfitting, we excluded MICAN, because this program was partially trained on the MALIDUP dataset^26^. The result shows that although US-align was not optimized to resemble manual alignments, it achieves a reasonable agreement to manual alignments with a F1-score 0.782, which is 3.3%, 10.3%, and 27.2% higher than those achieved by SPalign (0.757), Dali (0.709), and SSM (0.615), respectively. This probably reflects that the TM-score, which was designed to optimize the alignment accuracy and coverage simultaneously, has naturally captured the overall topological similarity of structures that is essential for the function and evolution of macromolecules.

### MSTA

MSTA matches multiple (3 or more) monomeric structures with similar topology into a single alignment matrix. To examine the ability of US-align for RNA MSTA, we collected a benchmark dataset by clustering the 637 structures from the RNA monomer alignment dataset used above by our in-house qTMclust algorithm (**Supplementary Text 4, Supplementary Fig. 5**) at a TM-score_RNA_ cutoff of 0.45. This resulted in 275, 39, and 31 clusters with 1, 2, and ≥ 3 chains, respectively. The 31 groups with ≥ 3 structures per group were used as the MSTA benchmark dataset in which multiple RNA alignments were performed within each cluster.

**Fig. 5** shows the average performance of US-align in comparison with four third-party programs^18, 19, 32, 33^, which were extended from RNA sequence and protein structure alignment tools (**Supplementary Text 5**). The comparison was based on a subset of 29 groups of RNAs for which all programs could generate results, since MUSTANG was not able to complete MSTA for two groups of long RNAs as explained by the **Supplementary Fig. 6** caption. The performance on the full set of all 31 RNA groups is shown in **Supplementary Fig. 6** and **Supplementary Table 6**. We note that it is not completely fair to pit US-align against LinearTurboFold and MAFFT-xinsi, as the latter two programs are extended from multiple sequence alignment (MSA) of RNA sequences and secondary structures rather than *bona fide* 3D structure alignment tools (**Supplementary Text 5**). Nonetheless, given the rareness of RNA MSTA programs, we include the programs in the comparison, which on one hand generates quite reasonable alignment results and one the other hand can help highlight the additional values that 3D structure information brings on top of MSA. As MSA programs, LinearTurboFold and MAFFT-xinsi tended to maximize the overlap among sequences, leading to very high coverages but with more than 4-times larger RMSDs and more than 11% lower TM-score_RNA_, compared to 3D structure alignments by US-align. US-align also outperformed the two MSTA programs (Matt and MUSTANG) by achieving 4.8% and 3.5% higher TM-score_RNA_ as well as 15.5% and 63.9% lower RMSD, respectively. Here, TM-score, RMSD, and coverage were all calculated from the pairwise alignments extracted from the MTSA. The result also shows that US-align was much faster than the control programs with the average time 15.0, 1650.3, 12.7 and 2.3 folds shorter than Matt, MUSTANG, LinearTurboFold, and MAFFT-xinsi, respectively.

**Fig. 5.**
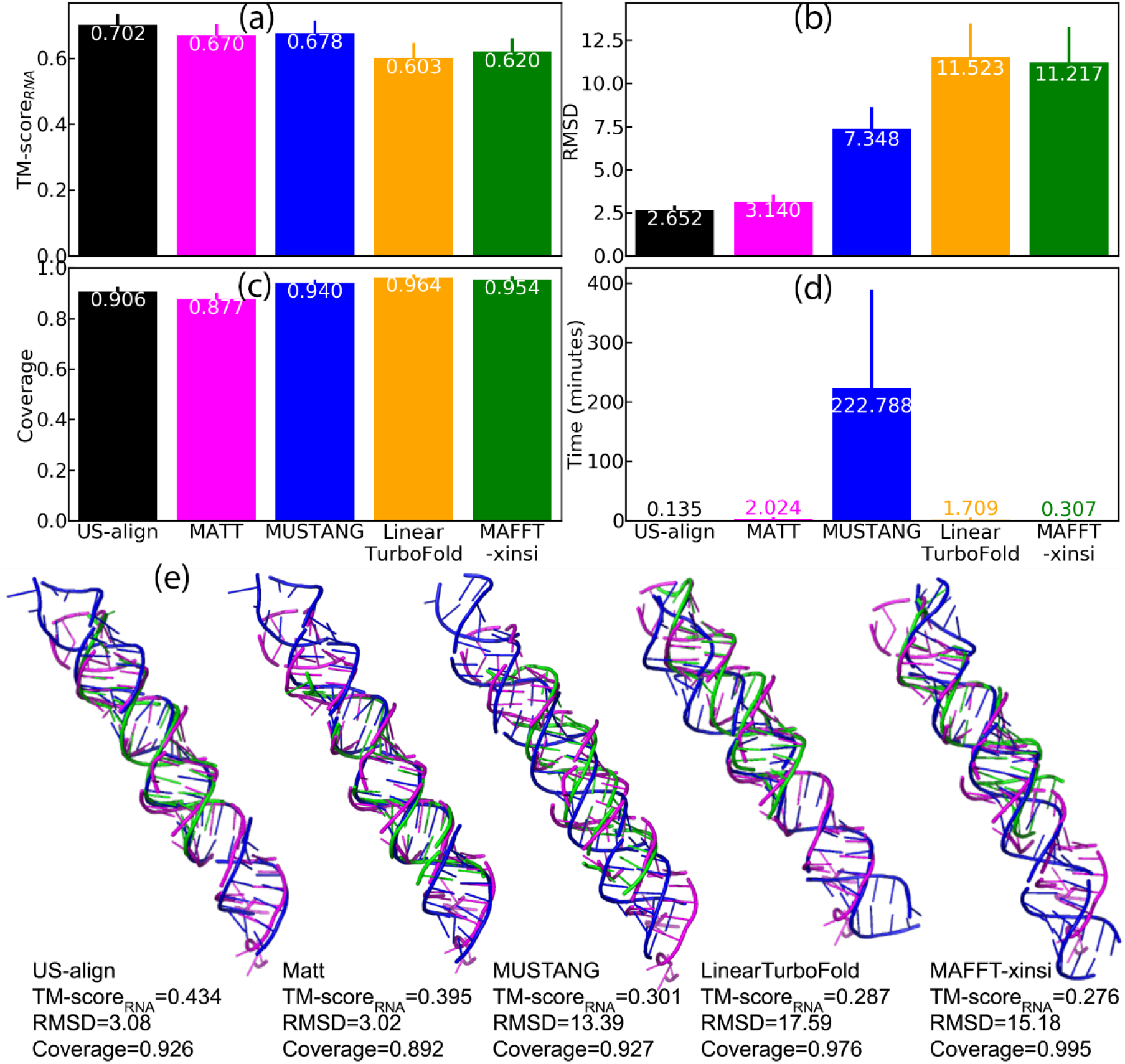
Multiple RNA alignment by US-align, Matt, MUSTANG, LinearTurboFold, and MAFFT-xinsi. **a-d**, Average performance measured by **a**, TM-score_RNA_, **b**, RMSD, **c**, alignment coverage, and **d**, running time. The length of an error bar represents the standard error of mean (SEM). Detailed per-target TM-score_RNA_, RMSD, and coverage information is shown in **Supplementary Fig. 6. e**, Superimposed structures derived from multiple alignment among three RNAs: PDB 6V5B chain D (78 nucleotides, magenta), PDB 2L3J chain B (71 nucleotides, blue), and PDB 2NUE chain C (46 nucleotides, green).

As a case study, **Fig. 5e** shows the MTSA for a group of three RNAs: a pri-miRNA (PDB 6V5B chain D), a pre-mRNA (PDB 2L3J chain B), and a dsRNA being processed by RNase III (PDB 2NUE chain C). Although all three structures have a simple topology (a single helix), LinearTurboFold, MAFFT-xinsi, and MUSTANG all failed to derive the correct correspondence between nucleotides of different structures, resulting in poor RMSD >13 Å. Both US-align and Matt created correct alignments with RMSD ∼3 Å, but US-align aligns more nucleotides and results in a higher coverage and TM-scrore_RNA_ (**Fig. 4e**).

In **Supplementary Fig. 7**, we further test the ability of US-align on protein MTSA in control with four state-of-the-art methods: PROMALS3D^34^, Matt^18^, MAMMOTH-mult^35^, and MUSTANG^19^. The benchmark dataset consists of 803 protein structures from 92 SCOPe fold families, where each fold family contains 3 to 42 structures which share the same fold but from different superfamilies^36^. Among the methods, US-align achieves the lowest pairwise RMSD (3.9 Å) with the second highest alignment coverage (68.7%). The average TM-score is 36.9-43.3% higher than other control methods, where the TM-score difference is statistically significant with a *p-*value below 1.3E-12 for all the comparisons (**Supplementary Table 7**). Meanwhile, the speed of US-align is on average 199.6, 24.6, 1.1, 30.7 times faster than PROMALS3D, Matt, MAMMOTH-mult, and MUSTANG, respectively.

### RNA-protein docking

Given the ability of US-align for both protein and nucleic acid structure alignments, we constructed a template-based RNA-protein docking pipeline by separately matching the query RNA and protein chains to a library of known RNA-protein complex structures, with the final models sorted by the root mean square of TM-scores of the RNA and protein structural alignments.

In **Fig. 6a-c**, we present a summary of performance of US-align on a set of 439 non-redundant RNA-protein complexes, in control with two state-of-the-art RNA-protein docking methods 3dRPC^37^ and PRIME^9^ which perform template-free and template-based docking, respectively (**Supplementary Text 6**). It was shown that US-align achieved a significantly lower median RNA RMSD than 3dRPC (by 15.5%) and PRIME (by 22.8%). If we define a successful case as those with RNA RMSD <10 Å, the success rate of US-align is 45.6% higher than 3dRPC and 13.8% higher than PRIME. Importantly, the average running time of US-align (19.89 min) is 28 times faster than 3dRPC (559.86 min) and 6 times faster than PRIME (118.49 min). In **Fig. 6d**, we present an illustrative example from the complex between a ribosomal protein and an mRNA (PDB ID 2VPL), where US-align created a model with a significantly lower RMSD (1.0 Å) than 3dRPC (29.3 Å) and PRIME (8.9 Å). Although PRIME and US-align recognized the same template (PDB ID 1MZP), the US-align model is much closer to the native structure due to more precise RNA and protein structure alignments.

**Fig. 6.**
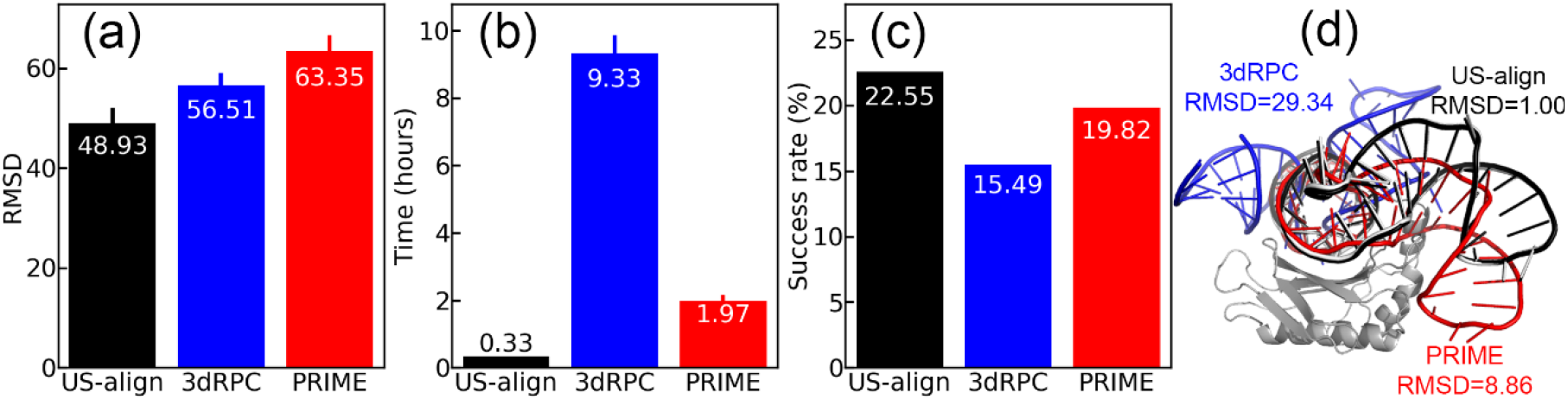
Overall performance of RNA-protein docking by US-align (black), 3dRPC (blue), and PRIME (red) in terms of **a**, median RMSD of docked RNA ligand, **b**, average running time, and **c**, success rate measured by the percentage of targets with a ligand RMSD<10 Å. The length of an error bar represents the standard error of mean (SEM). P-values are shown in **Supplementary Table 8. d**, Example of RNA-protein docking between ribosomal protein and mRNA (chain A and B, respectively, of PDB ID 2VPL, whose native structure is shown in grey).

## Conclusion

We developed US-align, a universal protocol for monomeric and oligomeric structural alignment of protein, RNA, and DNA molecules, built on the coupling of a uniform TM-score objective function and the heuristic iterative searching algorithm. Large-scale benchmarks show that US-align outperforms state-of-the-art programs in terms of both alignment accuracy and speed for a wide range of structural comparison tasks, including oligomeric structural alignment, RNA and protein MSTA, and template-based protein-RNA docking. Given the fundamental importance of structure comparisons in molecular biology, the high efficiency of a uniform structural alignment tool should significantly facilitate the related structural biology and function annotation studies across different types of biomolecules.

Despite the efficiency, US-align is essentially a tool for sequence-order dependent rigid structural alignments which may not be sufficient for some specific applications. For example, sequence-order independent (SOI) alignment is often preferred for comparing the binding pockets of ligand-receptor interactions in virtual screening studies. Meanwhile, flexible structure alignment may be needed for aligning multi-domain structures with alternative inter-domain orientations or for comparing multi-chain complexes with large conformational changes. Future developments will focus on extension of US-align for SOI and flexible alignments.

## Methods

### Monomeric Structure Alignment

To structurally align a pair of chains in US-align, it was necessary to derive the optimal alignment (i.e., the residue-level equivalence) between the two chains that maximizes the TM-score of structural superimpositions. For this, US-align starts with five sets of initial alignments:

1. Alignments from gapless sliding of one chain against another, and the alignment with the best TM-score was selected.
2. Alignment of the secondary structures of the two chains by Needleman-Wunsch (NW) dynamic programming^38^, using a gap penalty of -1, a match score of 1, and a mismatch score of 0.
3. Alignment based on NW dynamic programming, but the matching score is a half-half combination of secondary structure match and the residue-level TM-score calculated based on the superposition from Initial Alignment (1):

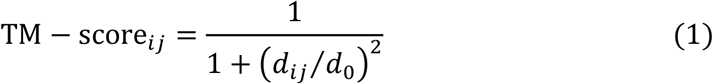

where *d*_*ij*_ is the distance between *i-*th on the first structure and *j-*th residue on the second structure.
4. Alignments based on the superimposition of fragments with length *L*_*min*_/2 and *L*_*min*_/3, where *L*_*min*_ is the minimal length between the two query chains. For saving time, a fragment is taken only every *n*_*jump*_ residues, where *n*_*jump*_ = min (45, *L*/3).
5. Alignment based on gapless sliding of all continuous fragments. For proteins, a fragment is ‘continuous’ if it has at least 4 residues and all C*α* -C*α* distances between adjacent residues are <4.25Å. For nucleic acids, any 4 nucleotides adjacent in the sequence are considers a piece of continuous fragments.

Each of the initial alignments is followed by a heuristic iteration alignment process, in which we first rotate the structures by TM-score rotation matrix based on the aligned residues in the initial alignment. Next, a new alignment is derived using NW dynamic programming, based on the residue-level matching score (Equation 1, calculated from the new superposition) with a gap penalty -0.6. The new alignment will result in a newer superposition which will be used to create a newer alignment. The process is repeated until convergence, where the structural alignment with the highest TM-score is returned. The overall procedure of monomeric structure alignment is illustrated by **Supplementary Fig. 8**.

### Oligomeric Structure Alignment

One challenge to oligomer complex alignment is chain equivalence assignment, i.e., finding the correct chain-level correspondence. In the simplest scenario of aligning two dimers, US-align needs two separate structural alignments: one for aligning chains A and B from dimer 1 to chains A and B in dimer 2, respectively; and another for aligning chains A and B from dimer 1 to chains B and A in dimer 2, respectively. More generally, when aligning an oligomer with *C*_1_ chains to another oligomer with *C*_2_ chains where *C*_1_ ≥ *C*_2_, US-align needs to determine the best chain assignment with the highest TM-score out of all *C*_1_!/(*C*_1_ − *C*_2_) ! possible chain assignment combinations. For example, alignment of a pair of octamers requires consideration of 8!/(8 − 8) ! = 40,320 possible chain assignments. This makes the exhaustive search approach, such as that used by MM-align^16^, extremely time consuming.

Therefore, when aligning oligomers with three or more chains, US-align employs a light-weighted chain assignment method. First, all-against-all chain-to-chain alignments are performed between all chains in oligomer 1 and all chains in oligomer 2 using fTM-align^39^, a fast version of TM-align. Compared to the standard TM-align, fTM-align decreases the number of iterations, thereby significantly reducing the required computing time while maintaining TM-scores highly correlated with those from standard TM-align^39^, especially for very large structures (**Supplementary Fig. 9**). The TM-scores from fTM-align are then used for initial chain assignment using the Enhanced Greedy Search (EGS)^40^ algorithm, by maximizing the sum of the TM-scores for all assigned chain pairs (**Supplementary Fig. 10**).

Once an initial chain assignment is decided, US-align will perform a TM-score superimposition of the two oligomers according to the inter-oligomer residue-level alignments generated in the previous step (**Supplementary Fig. 11**). Next, based on the TM-score superimposition matrix (Equation 1), a new optimal structure alignment will be obtained by a modified NW dynamics program which ignores the regions of unassigned chains (**Supplementary Fig. 12**). Given the new structural alignment, the chain-to-chain TM-scores will be computed for all inter-chain pairs and used by EGS to determine a new set of chain assignments (**Supplementary Fig. 10**), which will be returned to the last step for oligomer alignment iterations. This iteration will be repeated until convergence, where the structural alignment with the highest TM-score encountered during the iteration will be returned as the final structural alignment of the input oligomers (**Supplementary Fig. 11**).

### MSTA

To create a uniform alignment for multiple structures, US-align first performs all-against-all alignments among all input structures to obtain the pairwise TM-scores. Next, a structure-based guide tree is constructed based on the TM-scores using the extended unweighted pair group method with arithmetic mean (UPGMA) algorithm^41^ (**Supplementary Fig. 13**). Finally, the pairwise alignments from the first step are progressively merged into a single MSTA according to the branching order of the UPGMA tree. To merge an alignment with *M* structures to another alignment with *N* structures, NW dynamic programming is performed using a generalized version of the residue-level TM-score from Equation (1):

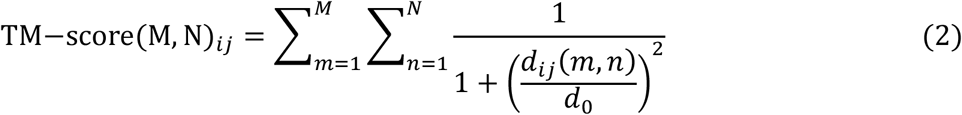

where *d*_*ij*_(*m, n*) is the distance of *i-*th residue position of the *m*-th structure in the first alignment and *j-*th residue position of the *n*-th structure from the second alignment after the superposition. The overall workflow of MSTA is illustrated by **Supplementary Fig. 14**.

### Template-based docking

To perform template-based docking, US-align implements a subroutine to align multiple query chains to one complex template. In this subroutine, each query chain is aligned to every chain of the complex template to calculate the TM-score. Based on the TM-scores, each query chain is then superimposed to one of the template chains so that no more than one query chain is assigned to the same template chain and the overall docking TM-score is maximized:

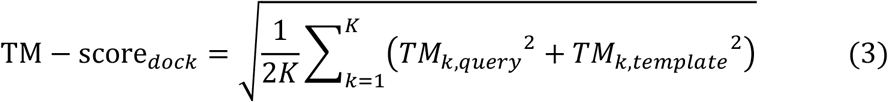

where *K* is the total number of query chains; *TM*_*k,qurey*_ and *TM*_*k,template*_ are the TM-score (or TM-score_RNA_) for aligning the *k*-th query chain, as normalized by the chain length of query and template, respectively.

When multiple templates were available, we ran US-align template-based docking on each complex template, and the template with the highest TM − score_*dock*_ was used to generate the final docking result.

## Data availability

All structure files and source code needed to reproduce this work are available at https://doi.org/10.6084/m9.figshare.16725745.

## Code availability

An online webserver and the standalone program of US-align are available at https://zhanggroup.org/US-align. The source code of US-align is also available at https://github.com/pylelab/USalign. The code was tested on Linux, Windows, and Mac OS, where no major difference in speed across different operating systems were found (**Supplementary Fig. 15**).

## Acknowledgements

We thank Sizhen Li from Dr. Liang Huang’s group at Oregon State University for assistances on LinearTurboFold. We thank Dr. Yang Cao for technical assistances in developing qTMclust. We thank Dr. Xiaoqiong Wei for insightful discussions. This work used the Extreme Science and Engineering Discovery Environment (XSEDE), which is supported by National Science Foundation grant number ACI1548562. C.Z. is a Howard Hughes Medical Institute postdoctoral fellow. A.M.P. is a Howard Hughes Medical Institute Investigator. This work is supported in part by the National Human Genome Research Institute (HG011868), National Institute of General Medical Sciences (GM136422, OD026825), the National Institute of Allergy and Infectious Diseases (AI134678), and the National Science Foundation (IIS1901191, DBI2030790, MTM2025426).

## Author contributions

Y.Z. conceived the study. C.Z., A.M.P. and Y.Z. designed the experiments. C.Z. developed the method. C.Z. and M.S. drafted the manuscript. All authors revised the manuscript and approved the final version.

## Ethics declarations

The authors declare no competing interests.

