## Supplementary for "US-align: Universal Structure Alignments of Proteins, Nucleic Acids, and Macromolecular Complexes"

### Supporting Information

#### Table of Contents

**Supporting Texts**

**Supplementary Text 1.** TM-score and TM-score$_{RNA}$.
**Supplementary Text 2.** Benchmark dataset for oligomeric structure alignment.
**Supplementary Text 3.** Q-score and Dali Z-score.
**Supplementary Text 4.** Structure-based clustering by qTMclust.
**Supplementary Text 5.** RNA multiple structure alignment (MTSA) programs.
**Supplementary Text 6.** Docking parameters and the benchmark dataset for RNA-protein docking.

**Supporting Figures**

**Supplementary Figure 1.** An example homotrimer structure alignment between a viral 5 F protein and the nuclear pore complex for which US-align underperforms MM-align.
**Supplementary Figure 2.** US-align heterooligomeric alignment between two intact protein-nucleic acid complexes.
**Supplementary Figure 3.** Performance of pairwise monomeric protein structure alignment.
**Supplementary Figure 4.** Agreement of manually curated pairwise monomeric protein alignment in the MALIDUP dataset and the automatic structure alignment.
**Supplementary Figure 5.** Schematic of structure clustering by qTMclust.
**Supplementary Figure 6.** Performance of multiple structure alignment for each group of RNA structures.
**Supplementary Figure 7.** Performance of protein multiple structure alignment.
**Supplementary Figure 8.** Flowchart for pairwise monomeric alignments by US-align.
**Supplementary Figure 9.** Performance of standard TM-align versus fTM-align implemented by US-align.
**Supplementary Figure 10.** Illustration of chain assignment by Enhanced Greedy Search (EGS).
**Supplementary Figure 11.** Flowchart for the alignment of two oligomers by US-align.
**Supplementary Figure 12.** Illustration of modified NW dynamic programming to align a pair of trimers with cross-chain alignment prevented.
**Supplementary Figure 13.** Illustration of guide-tree construction by UPGMA for MSTA.
**Supplementary Figure 14.** Workflow of multiple structure alignment by US-align.
**Supplementary Figure 15.** Average running time of US-align on different operating systems.

**Supporting Tables**

**Supplementary Table 1.** The average, standard error of mean (SEM), and *p*-values for TM-score, RMSD, and alignment coverage in oligomeric structure alignments.
**Supplementary Table 2.** The average, standard error of mean (SEM), and *p*-values for TM-score$_{RNA}$, RMSD, and alignment coverage in pairwise RNA structure alignment.
**Supplementary Table 3.** The average, standard error of mean (SEM), and p-values for TM-score, RMSD, and alignment coverage for pairwise protein structure alignment.
**Supplementary Table 4.** The average, standard error of mean (SEM), and *p*-values for Q-score (Q), Dali Z-score (DaliZ), and the number of pairs with TM-score≥0.5 for pairwise protein structure alignment.

**Supplementary Table 5.** The average, standard error of mean (SEM), and *p*-values for precision, recall, and F1-score for the agreement between manual pairwise protein alignment and automatic structure alignment.

**Supplementary Table 6.** The average, standard error of mean (SEM), and *p*-values for $\text{TM-score}_{\text{RNA}}$, RMSD, and alignment coverage in multiple RNA alignment.

**Supplementary Table 7.** The average, standard error of mean (SEM), and *p*-values for TM-score, RMSD, and alignment coverage in multiple protein alignment.

**Supplementary Table 8.** The average, median, standard error of mean (SEM), and *p*-values for RMSD of RNA-protein docking.

#### References

### Supporting Texts

#### Supplementary Text 1. TM-score and TM-score$_{RNA}$.

TM-score was originally proposed to quantify the similarity between a pair of aligned proteins[1]:

$$\text{TM}-\text{score} = \frac{1}{L}\sum_{i=1}^{L_{ali}} \frac{1}{1+(d_i/d_0)^2} \quad (1)$$

where $L$ is the sequence length of the target structure; $L_{ali}$ is the number of aligned residue pairs; $d_i$ is the distance between the Cα atoms of the $i$-th pair of aligned residues; and $d_0$ is a normalization factor to scale the residue distance:

$$d_0 = \begin{cases} 1.24\sqrt[3]{L-15} - 1.8, & \text{if } L > 21 \\ 0.5, & \text{otherwise} \end{cases} \quad (2)$$

TM-score was later extended to TM-score$_{RNA}$ for comparing nucleic acid structures[2], which uses the same formula of Eq. 1 but with a slightly different normalization factor $d_0$:

$$d_0 = \begin{cases} 0.6\cdot\sqrt{L-0.5} - 2.5, & \text{if } L \geq 30 \\ 0.7, & \text{if } 24 \leq L \leq 29 \\ 0.6, & \text{if } 20 \leq L \leq 23 \\ 0.5, & \text{if } 16 \leq L \leq 19 \\ 0.4, & \text{if } 12 \leq L \leq 15 \\ 0.3, & \text{otherwise} \end{cases} \quad (3)$$

Based on the extensive statistics of protein and RNA structure families[2, 3], it was found that TM-score≥0.5 or TM-score$_{RNA}$≥0.45 corresponds to a protein or RNA pair with similar global structural topology.

There is no closed-form solution to compute the superimposition (i.e., the rigid body rotation and translation of one structure towards another) that maximizes the TM-score given the residue-level correspondence between a pair of structures. Therefore, the superimposition for TM-score was numerically derived by a heuristic iteration process. Specially, to obtain the optimal TM-score superimposition between two sets of $L_{ali}$ aligned residues, we extracted all continuous fragments with fragment length being $L_{ali}$, $L_{ali}/2$, $L_{ali}/4$, …, 4, where each pair of fragments is superimposed to each other using the Kabsch algorithm[4] by minimizing the RMSD. Next, all the residue pairs with distance below $d_0$ were collected and superposed again using Kabsch matrix. This process was repeated till the rotation matrix converged, where the superposition with the highest TM-score was finally returned.

#### Supplementary Text 2. Benchmark dataset for oligomeric structure alignment

To benchmark the oligomeric structure alignment algorithms (US-align, MM-align, and MICAN), a dataset was collected from the PDB including protein complexes with 2 to 8 chains, total sequence lengths up to 5,000 residues, and pairwise sequence identity <30%. Due to the specific input format requirement of MM-align and MICAN, we excluded multi-model NMR structures as well as structures whose asymmetric unit and biological assembly have different chain arrangements, even though US-align could correctly parse these structures. This resulted in an initial dataset of 4,422 dimers, 610 trimers, 1,030 tetramers, 129 pentamers, 357 hexamers, 60 heptamers, and 134 octamers. We further reduced the number of dimers, trimers, tetramers, and hexamers by only selecting the top 200 structures with the best resolutions for each oligomer type so that the numbers of structures in different oligomeric states were comparable.

**Supplementary Text 3. Q-score and Dali Z-score.**

Apart from TM-score, RMSD and coverage, we also evaluate the accuracy of pairwise protein structure alignment in terms of Q-score and Dali Z-score. The Q-score is the objective function used by the SSM program:

$$Q = \frac{{L_{ali}}^2}{{L_3}^2\left(1 + \frac{1}{9 \cdot L_{ali}} \sum_{i=1}^{L_{ali}} {d_i}^2\right)} \quad (4)$$

where $L_3 = \sqrt{L_1 \cdot L_2}$ is the harmonic average of sequence lengths between structure 1 ($L_1$) and structure 2 ($L_2$); $L_{ali}$ is the number of aligned residue pairs and $d_i$ is the distance between the $i$-th aligned residue pairs, defined in the same way as in TM-score (**Supplementary Text 1** Equation 1).

Meanwhile, the Dali Z-score is the objective function used by the Dali program:

$$Z = -2 + \frac{2}{m(L_3)} \sum_{i=1}^{L_{ali}} \sum_{j=1}^{L_{ali}} \begin{cases} 0.2, & \text{if } i = j \\ \left(0.2 - \frac{2\left|d_{ij1} - d_{ij2}\right|}{d_{ij1} + d_{ij2}}\right) exp\left[\frac{\left(d_{ij1} + d_{ij2}\right)^2}{-1600}\right], & \text{if } i \neq j \end{cases} \quad (5)$$

where $d_{ij1}$ and $d_{ij2}$ are the intramolecular distances between the $i$-th and $j$-th aligned residues within structure 1 and within structure 2, respectively; $m(L_3)$ is a normalization factor for the sequence length:

$$m(L_3) = \begin{cases} L_3 - 189.809, & \text{if } L_3 > 400 \\ 7.9494 + 0.70852L_3 + 0.00025895{L_3}^2 - 0.0000019156{L_3}^3, & \text{if } L_3 \leq 400 \end{cases} \quad (6)$$

**Supplementary Text 4. Structure-based clustering by qTMclust.**

For benchmarking MSTA, we needed a dataset in which structurally similar RNAs were clustered into groups. To accomplish this, we developed Quick TM-score-based Clustering (qTMclust), a structure clustering program based on US-align.

Unlike existing structure clustering programs such as SPICKER[5], Calibur[6], and MaxCluster[7], which were developed for clustering different conformations of the same molecule, qTMclust clusters a set of structures for different RNAs or proteins. In addition to the difference in inputs, the goals of these clustering programs are also completely different. Previous structure clustering methods aimed to find the consensus structure corresponding to the best free energy conformation; to this end, adaptive density-based clustering implemented by SPICKER was proven to be effective. For this study, qTMclust aimed to classify all input structures into groups based on a pre-defined cutoff (e.g., TM-score≥0.5 or $\text{TM-score}_{\text{RNA}}$≥0.45). For this purpose, both adaptive density-based clustering and the commonly used k-means clustering method were inappropriate because they did not adhere to a predefined similarity cutoff. Instead, incremental clustering was found by previous studies [8] to be the most efficient approach in such scenarios.

In the qTMclust protocol, all input structures are first ranked in descending order of length. The first structure in the list is the first cluster representative. Each remaining structure in the list is then used as a query to align to representative structures of established clusters. If the query structure is similar to an existing representative, it is grouped to the same cluster; otherwise, it becomes the representative of a new cluster. This process is repeated until all input structures are assigned to one cluster. The clustering scheme is illustrated in **Supplementary Figure 5**. In this study, the clustering of 637 non-redundant RNA chains required 38 CPU minutes.

**Supplementary Text 5. RNA multiple structure alignment (MTSA) programs.**

Since no programs have been previously developed specifically for RNA MSTA, to benchmark the ability of US-align, we extended two types of third-party programs for RNA MSTA comparison. First, LinearTurboFold[9] and MAFFT-xinsi[10] are RNA multiple sequence alignment (MSA) methods that align both RNA sequences and secondary structures. Second, Matt[11] and MUSTANG[12] are two MSTA programs originally developed for protein alignments. Since Matt and MUSTANG do not use protein-specific features, a simple modification to allow the programs to read C3' atoms from RNAs instead of Cα atoms from proteins was sufficient to convert them from protein MSTA to RNA MSTA programs. All alignment programs were run with default parameters, except for LinearTurboFold, which was run with the "--b1 -1" option to handle alignment of sequences with significantly different lengths.

Since LinearTurboFold and MAFFT-xinsi generate alignments without superimposing structures, we need additional structural superimposition steps to derive the TM-score$_{RNA}$ and RMSD for benchmark purpose. Specifically, from an original MSA generated by either program, we extract all pairwise sequence alignments in FASTA format. The pairwise sequence alignments, together with their structures, are then input to the TM-score program to generate the superimposition of the corresponding RNA chain structures, from which TM-score$_{RNA}$ and RMSD values are calculated.

**Supplementary Text 6. Docking parameters and the benchmark dataset for RNA-protein docking.**

Docking by PRIME was performed using the following parameters:

```
-mode 15 -system 4
rmsd.I_rms_One_Atom_Within_Cutoff=5.0
rmsd.fnat_One_Atom_Contact_Cutoff=10.0
model_outfile=complex
compare2native=NO
sort_by=tmscore
out_model_pdb_number=1
```

Docking by 3dRPC was performed using the following parameters:

```
-mode 9 -system 9
RPDock.grid_step=1
RPDock.out_pdb=1
```

For both programs, we kept all docking parameters at their default value as recommended by their user manuals. The only exception was the option "out_model_pdb_number=1" for PRIME and "RPDock.out_pdb=1" for 3dRPC, which meant that only the first RNA-protein complex structure model would be output, as we only compared the first model among US-align, PRIME, and 3dRPC.

This benchmark was performed on the PRIME dataset of 439 non-redundant high-resolution RNA-protein complexes, which was used as the template library for both US-align and PRIME. On average, the TM-scores by US-align among the RNA-protein complexes, among the protein components, and among the RNA components are 0.753, 0.562, and 0.764, respectively. The average TM-score on RNA-protein complexes is not higher than that on RNA monomers, although it is indeed higher than both RNA and protein monomers in some cases as shown in **Figure 3** and **Supplementary Figure 2**. This is not unexpected, as a pair of RNAs that are structurally similar at the monomeric chain level can have different modes of interactions with proteins; this can result in a lower overall TM-score of complex when the alternative binding modes adopted by the RNA are different from the optimal orientation based on the RNA structure alone.

To ensure a fair comparison between template-based and template-free docking, for each query RNA-protein pair, all homologous templates sharing >30% protein sequence identities or >80% RNA sequence identities to the input were excluded from template-based docking. To further make the docking more challenging and realistic, the amino acid side chains in the input protein were reconstructed by FASPR[13] without the RNA partner in order to mimic the unbound docking experiments. As an RNA interact with a protein through its phosphate and ribose backbones rather than through its nucleobases, the nucleotide backbones in the input RNA were reconstructed by Rcrane.CLI[14] in the absence of its protein partner for the similar purpose.

#### Supporting Figures

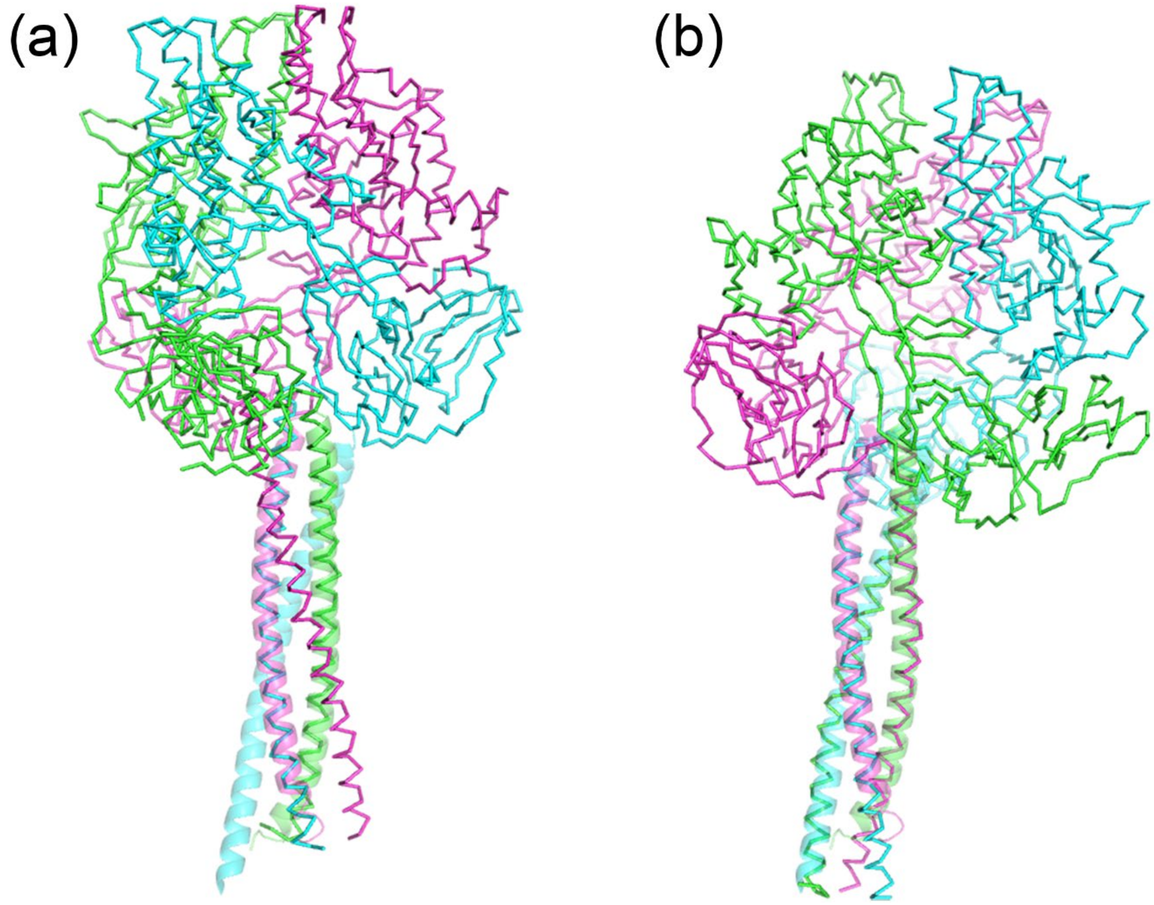


**Supplementary Figure 1.** An example homotrimer structure alignment between a viral 5 F protein (PDB 2B9B, ribbon) and the nuclear pore complex (PDB 3T97, semi-transparent cartoon) for which US-align underperforms MM-align. **a,** US-align (TM-score=0.525, initial chain assignment 2B98 Chain A, B, and C versus 3T97 Chain A, C, and B, respectively); **b,** MM-align (TM-score=0.892, initial chain assignment Chain A, B, and C versus 3T97 Chain A, C, and B, respectively). Chain A, B, and C are colored in green, cyan, and magenta, respectively.

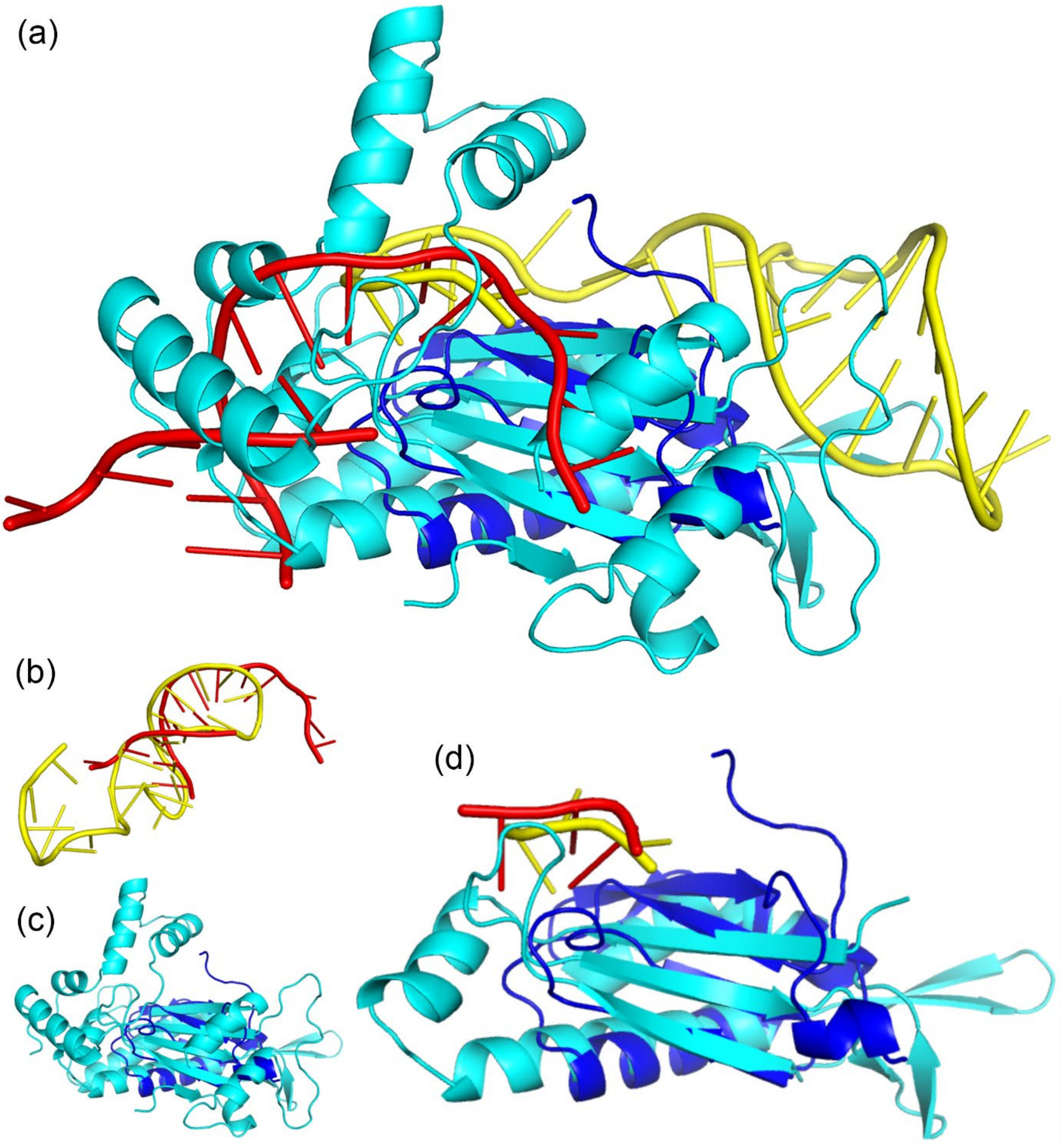


**Supplementary Figure 2.** US-align heterooligomeric alignment between two intact protein-nucleic acid complexes: spliceosomal protein U1A with its RNA substrate (PDB 1URN; chain A and P in blue and red, respectively) and relaxase TrwC with its DNA substrate (PDB 1OMH; chain A and B in cyan and yellow, respectively). **a,** Although the first structure was a protein-RNA complex and the second structure was a protein-DNA complex, US-align identified the similar region between the two structures at TM-score=0.467. **b-c,** On the other hand, the monomeric alignments for the proteins and for the RNA/DNA have only **b,** TM-score=0.301 and **c,** $\text{TM-score}_{\text{RNA}}$=0.157, respectively. **d,** Part of the reason for the difference in TM-score was that the proteins in both complexes bind to their nucleic acid substrate at a similar position as shown in this panel, where the N- and C-termini (residues 1-79 or residues 198-293) of 1OMH chain A are hidden as they were not aligned to 1URN. This panel also hid nucleotides that are not aligned in both structures.

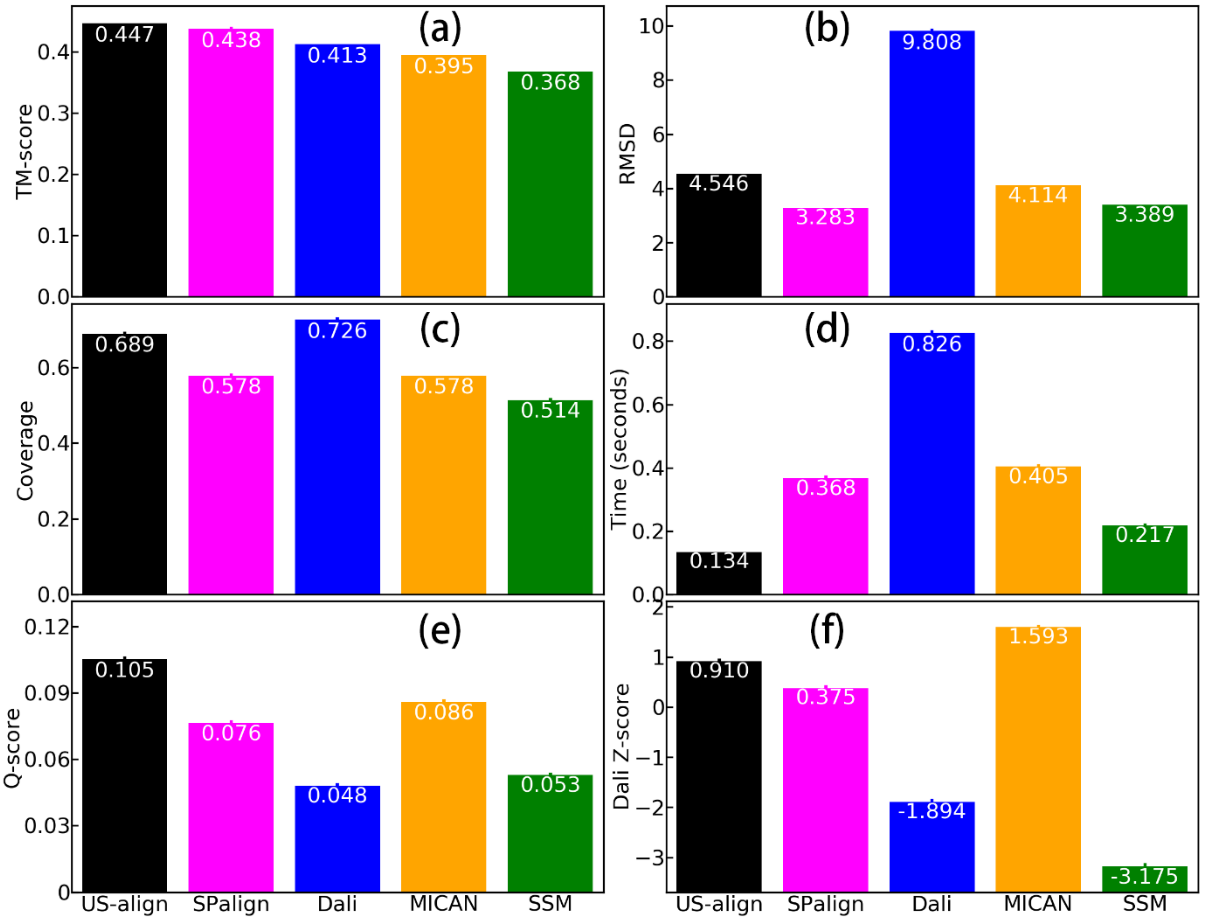


**Supplementary Figure 3.** Performance of pairwise monomeric protein structure alignment by US-align, SPalign[15], SSM[16], MICAN[17] and Dali[18]. The performance was measured by average **a**, TM-score, **b**, RMSD, **c**, alignment coverage, **d**, running time, **e**, Q-score and **f**, Dali Z-score. The dataset for this benchmark was generated by three steps. First, protein sequences for the ASTRAL set were downloaded from the SCOPe database[19] version 2.06. Second, the sequences were clustered by CD-HIT[8] at 30% sequence identity cutoff to remove redundant proteins, resulting in 9,896 representative sequences. Third, among these 9,896 structures, a subset of 1,000 structures with the best resolutions were included in the final dataset, leading to 1,000*999/2=499,500 pairs of chains. Since SSM and Dali cannot generate results for 3.4% and 93.5% of the pairs, this figure shows the results for the common set of 31,951 chain pairs for which all methods could generate alignment results. Error bars on top of most bars were not visible because the standard error of mean (SEM) values for all metrics were close to zero.

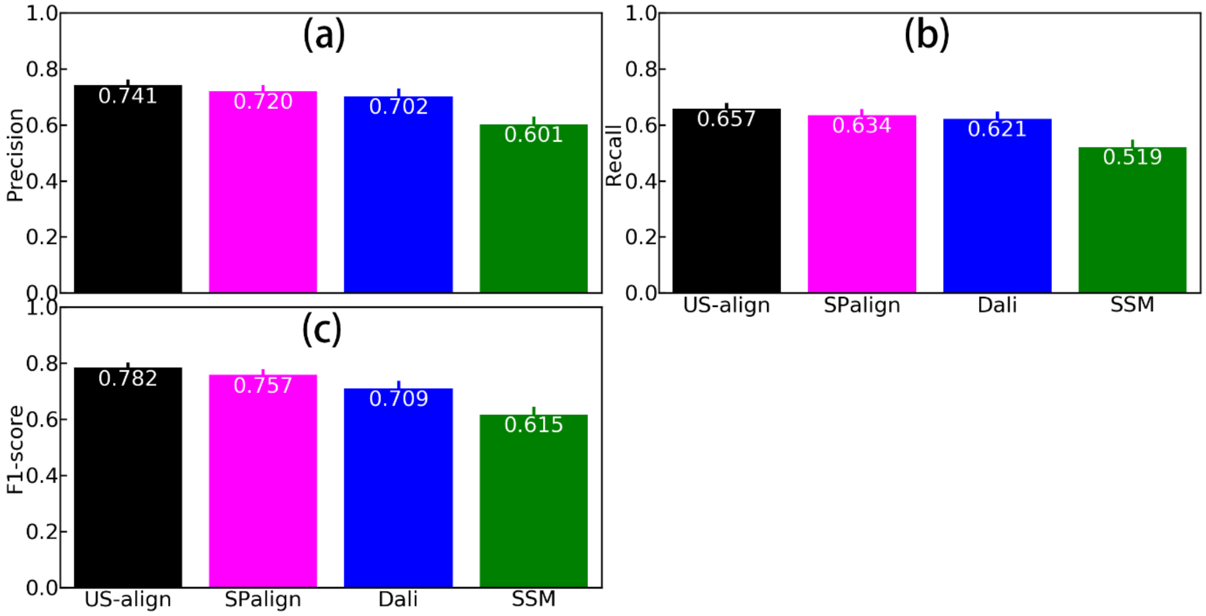


**Supplementary Figure 4.** Agreement of manually curated pairwise monomeric protein alignment in the MALIDUP[20] dataset and the automatic structure alignment by US-align, SPalign[15], SSM[16], MICAN[17], and Dali[18]. The performance was measured by **a**, precision (the number of aligned residue pairs consistent between manual and automatic alignment divided by the total number of automatically aligned residue pairs), **b**, recall (the number of aligned residue pairs consistent between manual and automatic alignment divided by the total number of manually aligned residue pairs), and **c**, F1-score (harmonic average between precision and recall). Error bars represent the standard error of mean (SEM) values across all 241 protein structure pairs in the MALIDUP dataset.

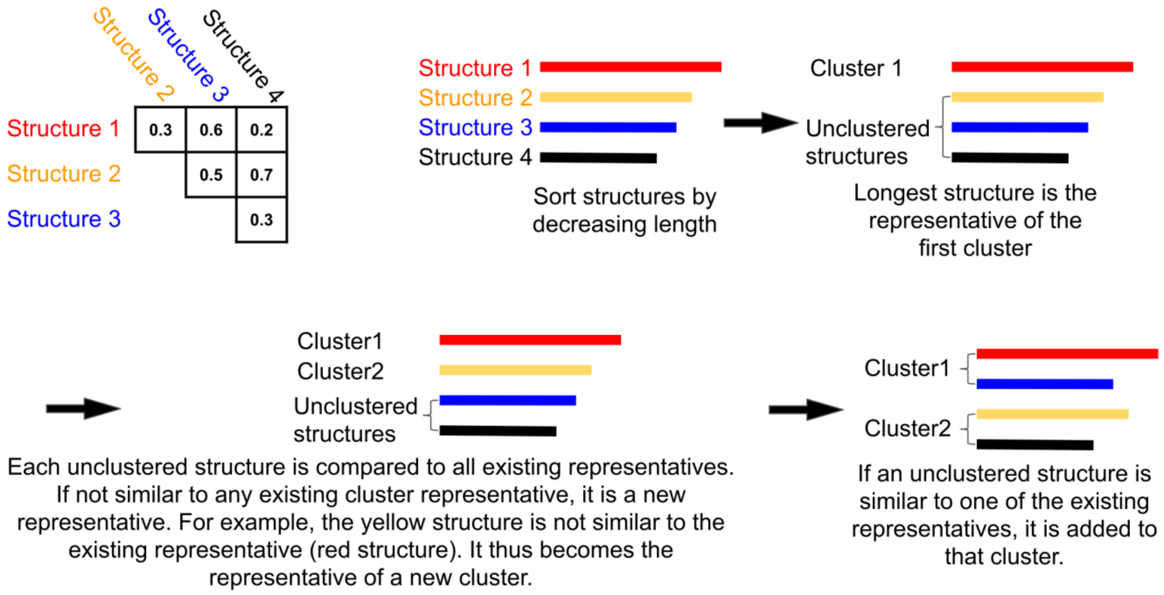


**Supplementary Figure 5.** Schematic of structure clustering by qTMclust. The upper left inset shows the TM-score matrix between different input structures, although in practice, it is not necessary to calculate the TM-score between non-representative members from different clusters (e.g., Structure 3 in blue versus Structure 4 in black in this illustration). In this study, we used the TM-score normalized by the longer structure in the alignment because we wanted to prevent the grouping of structures with too large differences in lengths into the same cluster. Nonetheless, qTMclust provided options to use the TM-score normalized by the shorter structure or by the average of the two TM-scores.

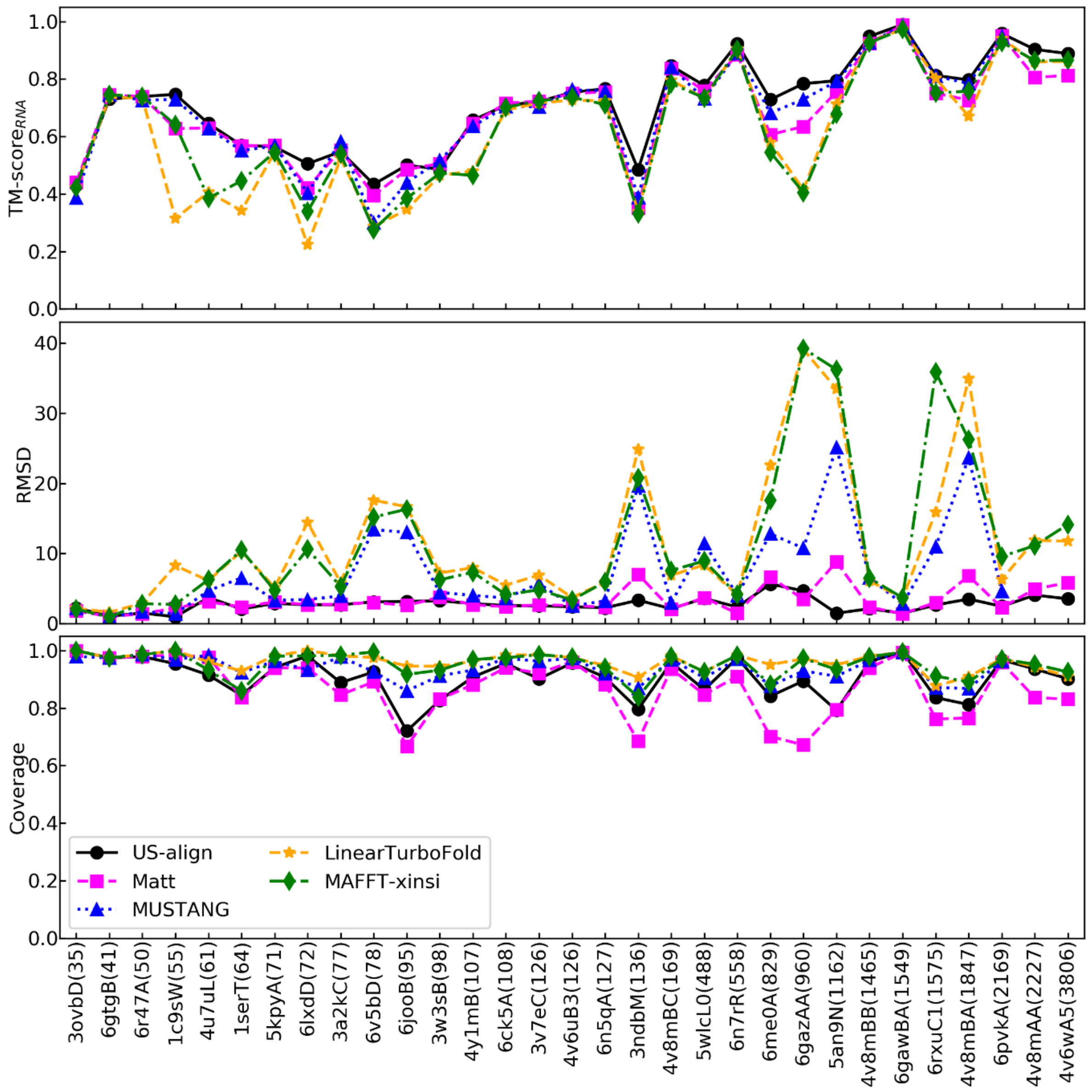


**Supplementary Figure 6.** Performance (average TM-score$_{RNA}$, RMSD, and alignment coverage on *y*-axis) of multiple structure alignment for each group of RNA structures (*x*-axis). RNA groups are ranked in ascending order of the length of the longest RNA in each group. For example, 3ovbD(35) on the *x*-axis represents a group of RNAs whose longest RNA is PDB 3OVB chain D with 35 nucleotides. MUSTANG results are unavailable for group 4v8mAA(2227) and 4v6wA5(3806) because their MSTA requires >30 days, which is the maximum running time on the Yale FARNAM supercomputer cluster used for this benchmark.

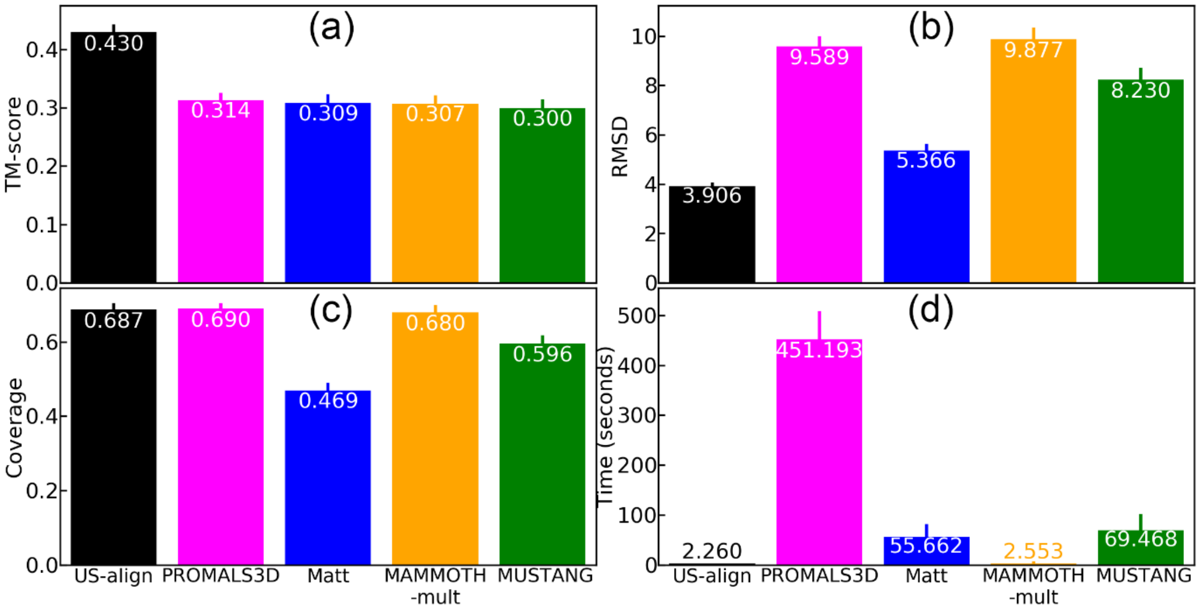


**Supplementary Figure 7.** Performance of protein multiple structure alignment by US-align, PROMALS3D[21], Matt[11], MAMMOTH-mult[22], and MUSTANG[12]. The performance was measured by average **a**, TM-score, **b**, RMSD, **c**, alignment coverage, and **d**, running time. The dataset for this benchmark was generated by three steps. First, protein sequences ranging from 30 to 1,000 residues for the ASTRAL set were downloaded from the SCOPe database[19] version 2.06. Second, for each SCOPe superfamilies, we only kept the structure with the best resolution. Third, out of the 1,221 SCOPe folds, we only kept the SCOPe folds with ≥3 superfamilies. This procedure resulted in 803 structures from 92 SCOPe folds, where each fold had 3 to 42 structures. Error bars show the standard error of mean (SEM) values.

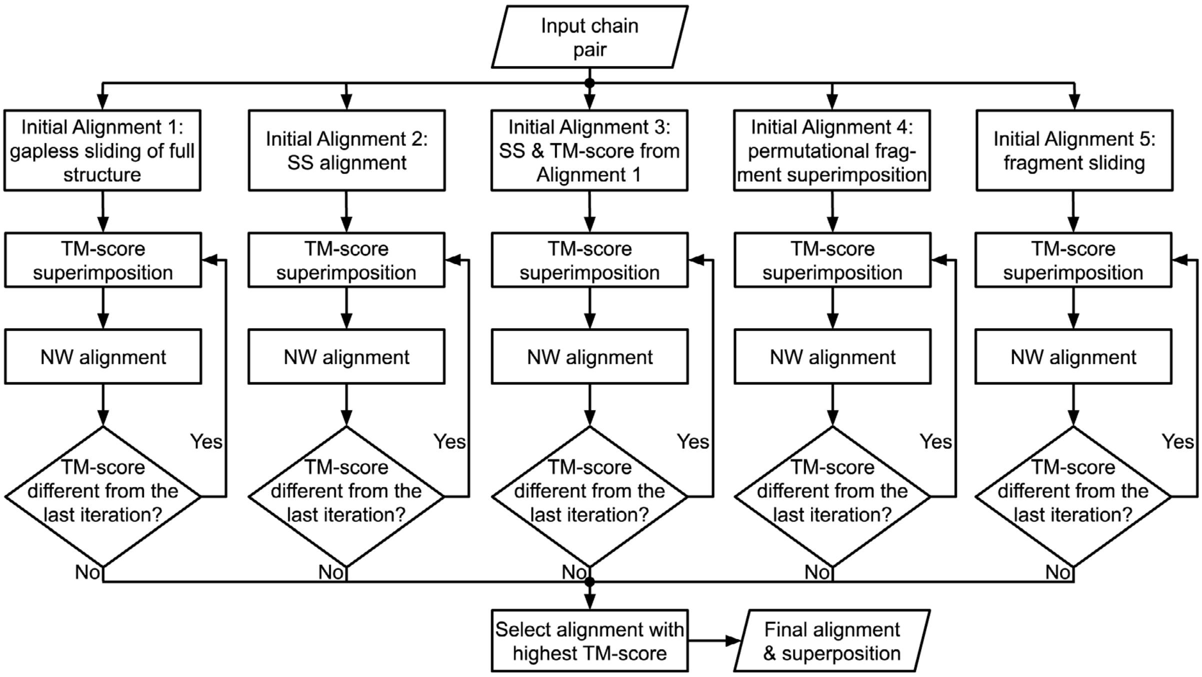


**Supplementary Figure 8.** Flowchart for pairwise monomeric alignments by US-align. Secondary structure is abbreviated as SS, and Needleman-Wunsch global alignment is abbreviated as NW.

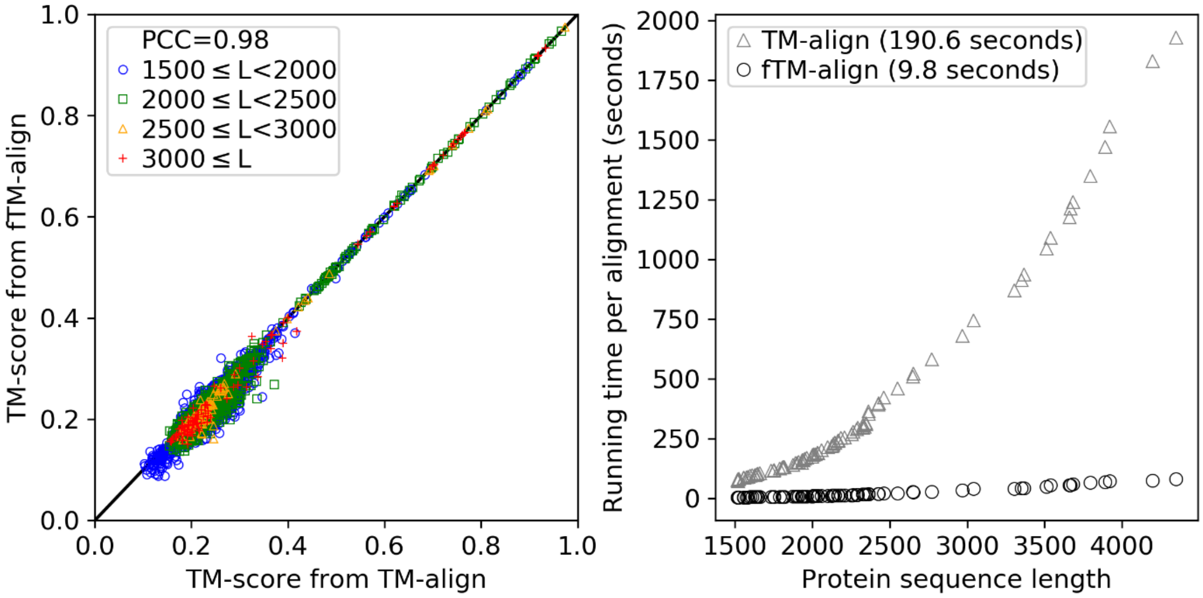


**Supplementary Figure 9.** Performance of standard TM-align versus fTM-align implemented by US-align for proteins with different lengths. PCC stands for Pearson Correlation Coefficient.

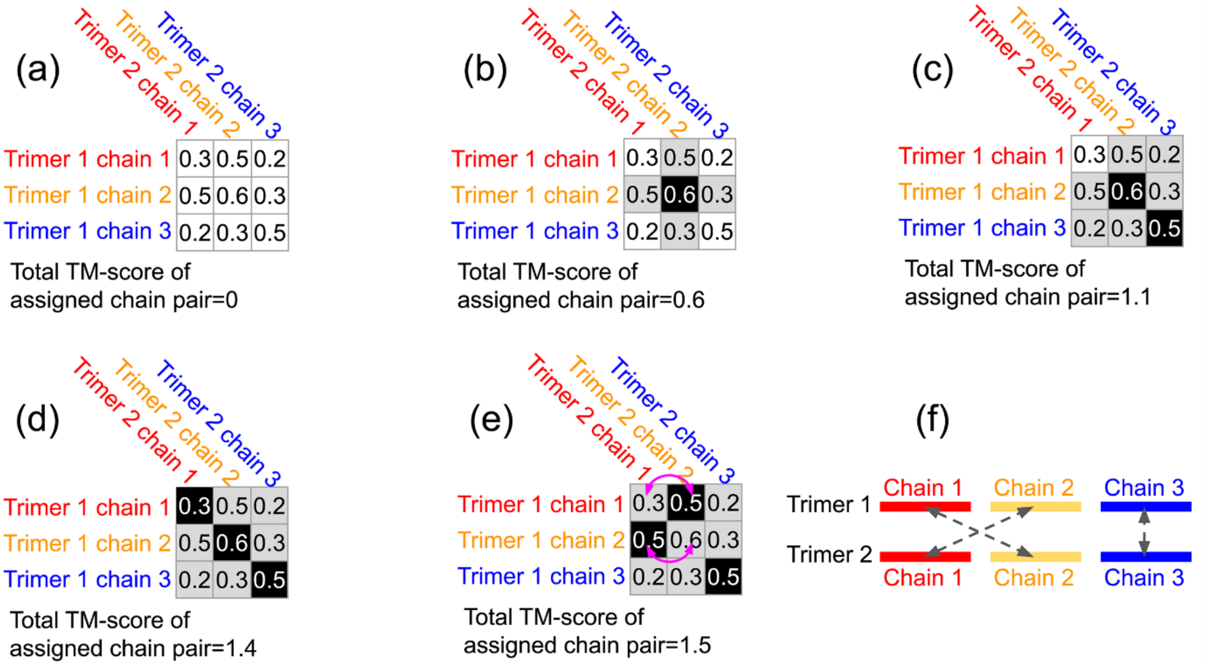


**Supplementary Figure 10.** Illustration of chain assignment by Enhanced Greedy Search (EGS) for a pair of trimers. **a,** All-against-all chain-to-chain TM-scores were computed. The TM-scores can be represented as a 3 by 3 matrix, where each row corresponds to a chain in trimer 1 and each column corresponds to a chain in trimer 2. **b,** The chain pair with the highest TM-score (black cell in the middle) was selected as the first assigned pair. Other cells in the same row or the same column (grey cells) were considered invalid assignments. **c,** The chain pair with the highest TM-score among the remaining cells was selected as the next assigned chain pair (black cell at lower right). Other cells in the same row or column were marked as invalid (grey). **d,** Step **c** was repeated until no more assignments could be made. **e,** For every two assigned chain pairs, the chain assignments were swapped (double arrows) if the swapping led to higher total TM-score. The swapping was repeated until no swap was possible. **f,** The final chain assignment is indicated by the dashed double arrows.

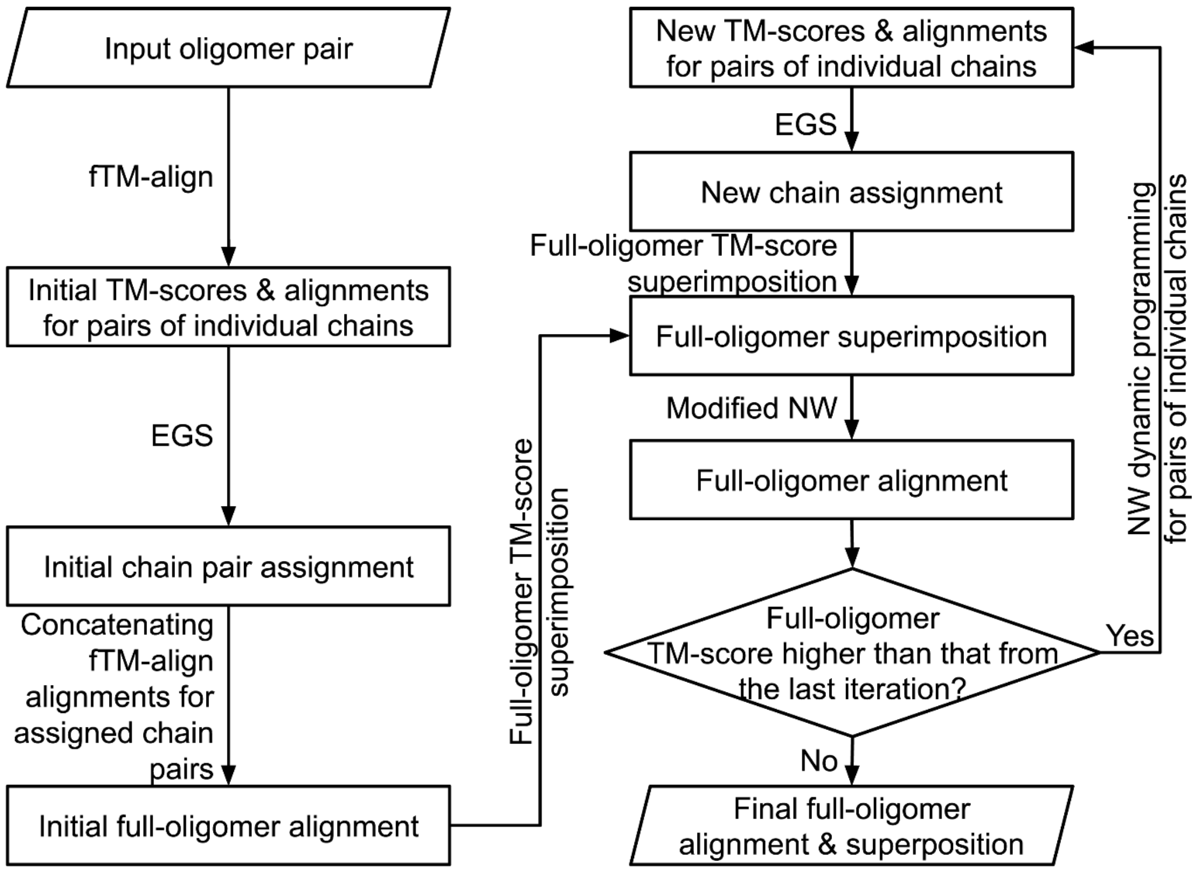


**Supplementary Figure 11.** Flowchart for the oligomer structural alignments by US-align. EGS is short for Enhanced Greedy Search (**Supplementary Figure 10**). fTM-align, which is a fast version of TM-align, is used to perform initial all-against-all chain-to-chain alignment between all chains in oligomer 1 and all chains in oligomer 2. Modified NW refers to the modified Needleman-Wunsch (NW) dynamic programming with cross-chain alignment prevented (**Supplementary Figure 12**).

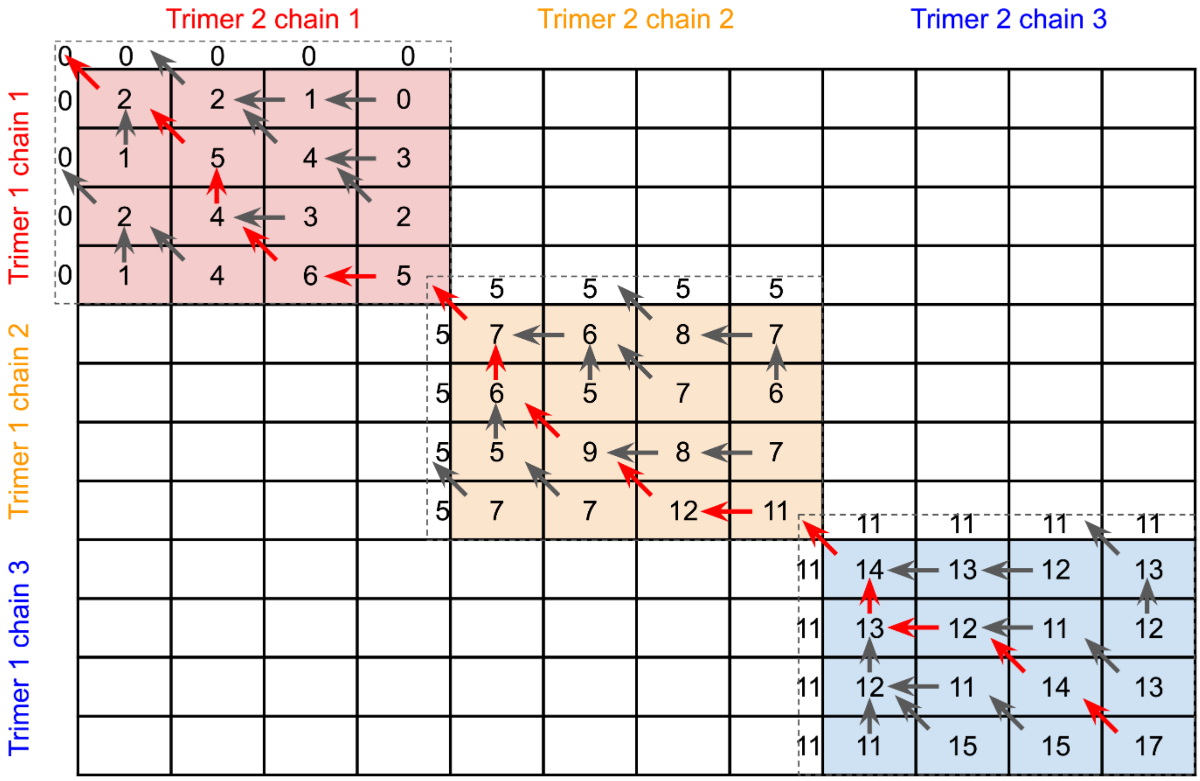


**Supplementary Figure 12.** Illustration of the modified Needleman-Wunsch (NW) dynamic programming to align a pair of trimers with cross-chain alignment prevented. In the dynamic programming matrix, only the color-shared regions corresponding to chain pairs assigned by Enhanced Greedy Search (EGS, **Supplementary Figure 10**) are filled up, while the remaining white regions for cross-chain alignments are ignored. In this example, chains 1, 2 and 3 from trimer 1 are assigned to chains 1, 2, and 3 from trimer 2, respectively. The dashed lines represent a pseudo-column/row which assumes the value in the last cell of the preceding block. The values of the pseudo-column/row (5 and 11 in this example) are used as starting score of the next block corresponding to the next chain of both complexes. The red arrows indicate the traceback path.

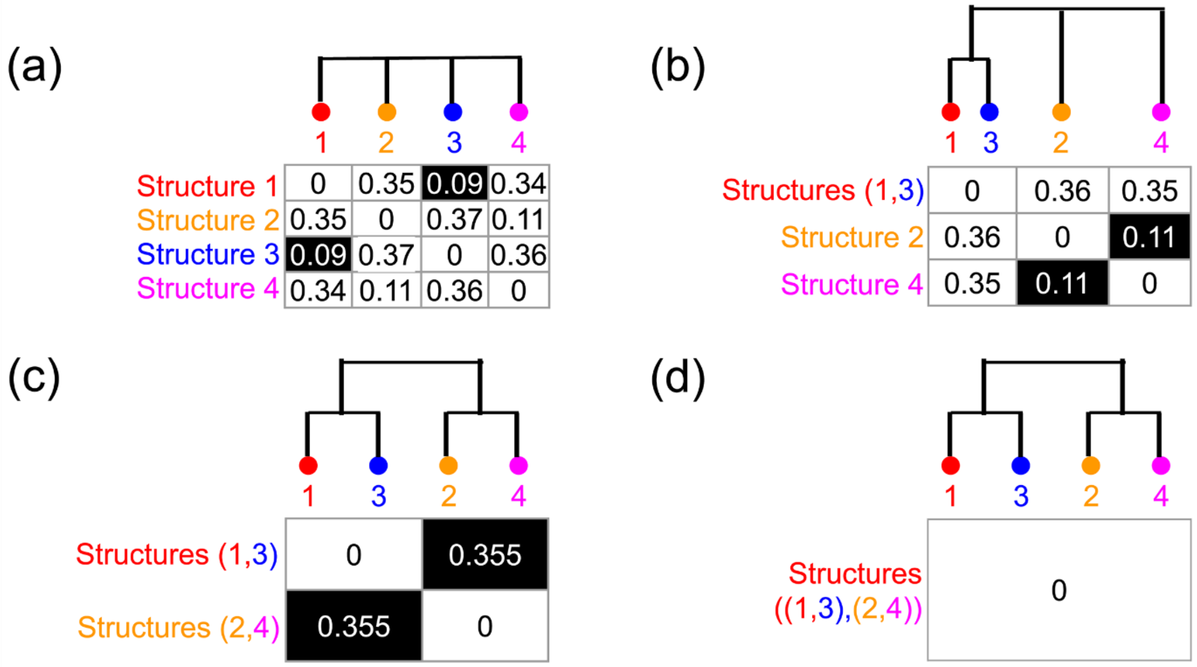


**Supplementary Figure 13.** Illustration of guide-tree construction by unweighted pair group method with arithmetic mean (UPGMA) for multiple structure alignment (MSTA) of four structures. **a,** Calculate the distance matrix from the all-against-all pairwise structure alignments, where the distance between two structures equals one minus the TM-score. Identify the structure pair with the smallest distance (black cell for Structure 1 and 3) among all pairs in the matrix, ignoring the distances on the matrix diagonal. **b,** Merge the pair with the smallest distance identified in the previous step into a single group. Recalculate the distance matrix. Here, the distance $d_{A,B}$ between Groups $A$ and $B$ with $|A|$ and $|B|$ structures, respectively, is calculated as:

$$d_{A,B} = \frac{1}{|A| \cdot |B|} \sum_{a=1}^{|A|} \sum_{b=1}^{|B|} d_{a,b}$$

where $d_{a,b}$ is the distance between structure $a$ and $b$ from Groups A and B, respectively. From the new distance matrix, find the structure group pair with the smallest distance (black cell for Structure 2 and 4). **c-d,** repeat the steps until all structure groups are merged.

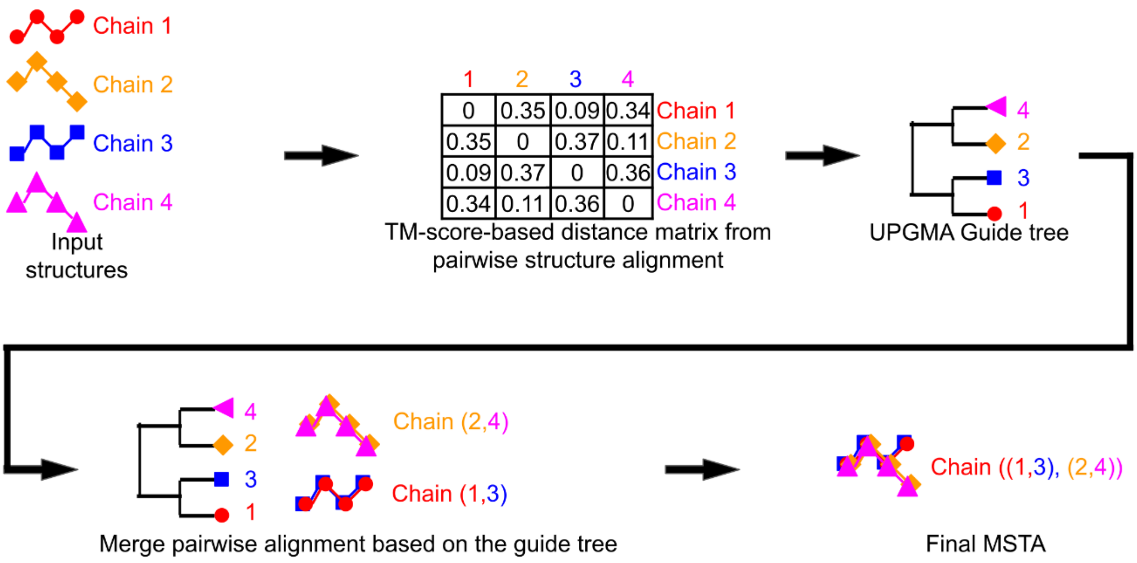


**Supplementary Figure 14.** Workflow of multiple structure alignment (MSTA) by US-align.

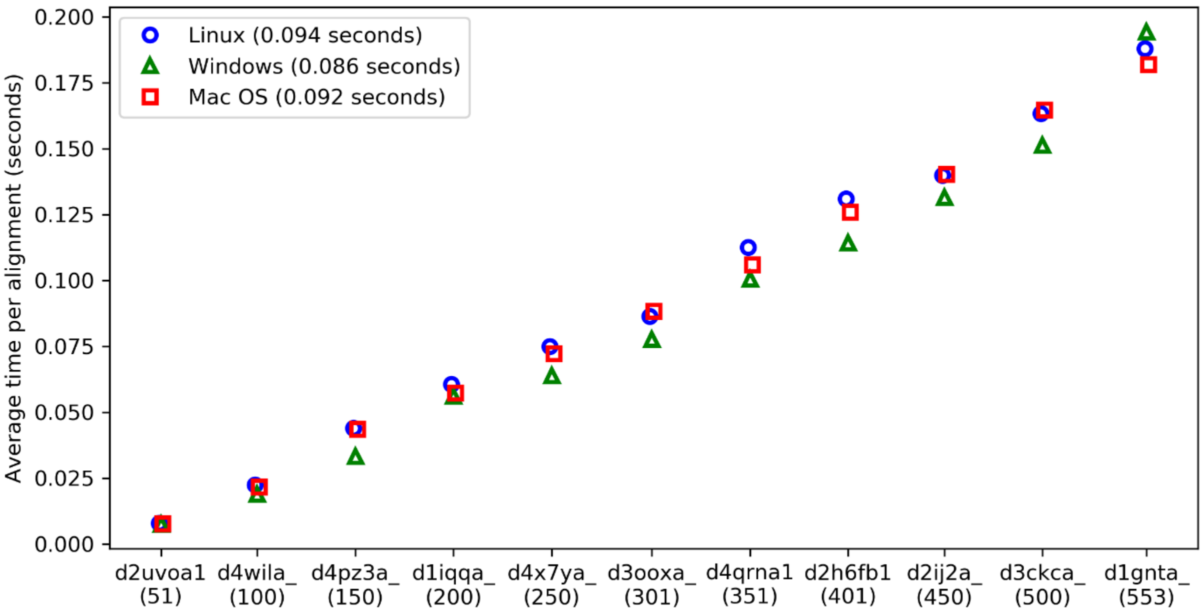


**Supplementary Figure 15.** Average running time of US-align on different operating systems (Ubuntu Linux 18.04, Windows 10, Mac OS 11) for 12 structures of different length (*x*-axis parentheses). These 12 proteins were selected by length from the 1,000 SCOPe structures in **Figure S3**. Parentheses in the upper left box shows the average running time across all structures. The average time is obtained by running US-align between each of the 12 selected proteins against all 1,000 SCOPe structures collected in **Supplementary Figure 3**. All source code compilation was done on 64-bit operating system by the g++ compiler using the "-O3 -ffast-math" option to obtain the highest level of speed optimization.

### Supporting Tables

**Supplementary Table 1.** The average, standard error of mean (SEM), and p-values for TM-score (TM), RMSD (RMS), and alignment coverage (COV) in oligomeric structure alignments by US-align, MM-align, and MICAN.

| Type | Methods | TM mean | TM SEM | TM *p*-value | RMS mean | RMS SEM | RMS *p*-value | COV mean | COV SEM | COV *p*-value |
|---|---|---|---|---|---|---|---|---|---|---|
| Dimer | US-align | 0.256 | 0.0004 | * | 7.13 | 0.01 | * | 0.418 | 0.001 | * |
| | MM-align | 0.250 | 0.0004 | 2.89E-159 | 7.20 | 0.01 | 8.12E-75 | 0.408 | 0.001 | 4.26E-147 |
| | MICAN | 0.207 | 0.0005 | <1E-303 | 5.44 | 0.01 | <1E-303 | 0.296 | 0.001 | <1E-303 |
| Trimer | US-align | 0.252 | 0.0005 | * | 7.38 | 0.01 | * | 0.411 | 0.001 | * |
| | MM-align | 0.249 | 0.0005 | 4.57E-40 | 7.43 | 0.01 | 7.63E-44 | 0.404 | 0.001 | 1.46E-48 |
| | MICAN | 0.216 | 0.0006 | <1E-303 | 5.71 | 0.00 | <1E-303 | 0.310 | 0.001 | <1E-303 |
| Tetramer | US-align | 0.225 | 0.0004 | * | 7.94 | 0.01 | * | 0.362 | 0.001 | * |
| | MM-align | 0.216 | 0.0004 | 1.19E-214 | 7.95 | 0.01 | 5.72E-3 | 0.347 | 0.001 | 1.34E-265 |
| | MICAN | 0.209 | 0.0005 | <1E-303 | 5.86 | 0.00 | <1E-303 | 0.294 | 0.001 | <1E-303 |
| Pentamer | US-align | 0.257 | 0.0009 | * | 8.06 | 0.02 | * | 0.416 | 0.001 | * |
| | MM-align | 0.233 | 0.0007 | <1E-303 | 8.07 | 0.02 | 1.12E-1 | 0.375 | 0.001 | <1E-303 |
| | MICAN | 0.233 | 0.0009 | 1.08E-202 | 5.97 | 0.01 | <1E-303 | 0.330 | 0.001 | <1E-303 |
| Hexamer | US-align | 0.229 | 0.0005 | * | 9.20 | 0.01 | * | 0.364 | 0.001 | * |
| | MM-align | 0.206 | 0.0004 | <1E-303 | 9.16 | 0.01 | 1.03E-20 | 0.324 | 0.001 | <1E-303 |
| | MICAN | 0.201 | 0.0005 | <1E-303 | 6.24 | 0.00 | <1E-303 | 0.278 | 0.001 | <1E-303 |
| Heptamer | US-align | 0.252 | 0.0025 | * | 8.82 | 0.04 | * | 0.398 | 0.003 | * |
| | MM-align | 0.222 | 0.0019 | 1.34E-67 | 8.81 | 0.04 | 8.12E-1 | 0.349 | 0.003 | 1.24E-83 |
| | MICAN | 0.224 | 0.0023 | 7.21E-54 | 6.13 | 0.01 | <1E-303 | 0.314 | 0.003 | 9.79E-164 |
| Octamer | US-align | 0.230 | 0.0006 | * | 9.40 | 0.02 | * | 0.366 | 0.001 | * |
| | MM-align | 0.191 | 0.0004 | <1E-303 | 9.32 | 0.01 | 1.44E-32 | 0.300 | 0.001 | <1E-303 |
| | MICAN | 0.216 | 0.0007 | 2.63E-106 | 6.33 | 0.00 | <1E-303 | 0.300 | 0.001 | <1E-303 |

* The *p*-values are from paired Student's t-tests against US-align. The *p*-values for some comparisons are too close to zero to be computed under IEEE double precision floats; these p-values are marked as "<1E-303" in the table.

**Supplementary Table 2.** The average, standard error of mean (SEM), and *p*-values for TM-score$_{RNA}$ (TM), RMSD (RMS), and alignment coverage (COV) for pairwise RNA structure alignment by US-align, RMalign, STAR3D, ARTS, and Rclick.

| Methods | TM mean | TM SEM | TM *p*-value | RMS mean | RMS SEM | RMS *p*-value | COV mean | COV SEM | COV *p*-value |
|---|---|---|---|---|---|---|---|---|---|
| US-align | 0.273 | 7.11E-7 | * | 3.805 | 8.15E-6 | * | 0.591 | 1.08e-6 | * |
| RMalign | 0.258 | 7.35E-7 | <1E-303 | 3.907 | 7.70E-6 | <1E-303 | 0.588 | 1.10E-6 | 3.05E-47 |
| STAR3D | 0.214 | 7.38E-7 | <1E-303 | 3.705 | 7.46E-6 | <1E-303 | 0.512 | 1.18E-6 | <1E-303 |
| ARTS | 0.203 | 7.45E-7 | <1E-303 | 3.818 | 8.15E-6 | 2.76E-10 | 0.442 | 1.12E-6 | <1E-303 |
| Rclick | 0.197 | 8.07E-7 | <1E-303 | 4.077 | 9.14E-6 | <1E-303 | 0.450 | 1.24E-6 | <1E-303 |

* The *p*-values are from paired Student's t-tests against US-align. The *p*-values for some comparisons are too close to zero to be computed under IEEE double precision floats; these p-values are marked as "<1E-303" in the table.

**Supplementary Table 3.** The average, standard error of mean (SEM), and *p*-values for TM-score (TM), RMSD (RMS), and alignment coverage (COV) for pairwise protein structure alignment by US-align, SPalign, SSM, MICAN, and Dali on the non-redundant SCOPe dataset.

| Methods | TM mean | TM SEM | TM *p*-value | RMS mean | RMS SEM | RMS *p*-value | COV mean | COV SEM | COV *p*-value |
|---|---|---|---|---|---|---|---|---|---|
| US-align | 0.447 | 5.70E-4 | * | 4.546 | 5.48E-3 | * | 0.689 | 6.70E-4 | * |
| SPalign | 0.438 | 5.91E-4 | 3.40E-26 | 3.283 | 2.44E-3 | <1E-303 | 0.578 | 8.19E-4 | <1E-303 |
| Dali | 0.413 | 6.38E-4 | <1E-303 | 9.808 | 2.46E-2 | <1E-303 | 0.726 | 8.70E-4 | 1.58E-246 |
| MICAN | 0.395 | 6.09E-4 | <1E-303 | 4.114 | 4.09E-3 | <1E-303 | 0.578 | 8.77E-4 | <1E-303 |
| SSM | 0.368 | 5.92E-4 | <1E-303 | 3.389 | 4.14E-3 | <1E-303 | 0.514 | 8.16E-4 | <1E-303 |

* The *p*-values are from paired Student's t-tests against US-align. The *p*-values for some comparisons are too close to zero to be computed under IEEE double precision floats; these p-values are marked as "<1E-303" in the table.

**Supplementary Table 4.** The average, standard error of mean (SEM), and *p*-values for Q-score (Q), Dali Z-score (DaliZ), and the number of pairs with TM-score≥0.5 for pairwise protein structure alignment by US-align, SPalign, SSM, MICAN, and Dali on the non-redundant SCOPe dataset.

| Methods | Q mean | Q SEM | Q *p*-value | DaliZ mean | DaliZ SEM | DaliZ *p*-value | TM-score ≥0.5 |
|---|---|---|---|---|---|---|---|
| US-align | 0.105 | 4.23E-4 | * | 0.910 | 2.22E-2 | * | 8119 |
| SPalign | 0.076 | 4.58E-4 | <1E -303 | 0.375 | 2.60E-2 | 7.48E-55 | 7661 |
| Dali | 0.048 | 3.82E-4 | <1E-303 | -1.894 | 3.26E-2 | <1E-303 | 6050 |
| MICAN | 0.086 | 3.94E-4 | 5.93E-242 | 1.593 | 1.81E-2 | 9.55E-125 | 4720 |
| SSM | 0.053 | 3.79E-4 | <1E-303 | -3.175 | 3.39E-2 | <1E-303 | 3268 |

* The *p*-values are from paired Student's t-tests against US-align. The *p*-values for some comparisons are too close to zero to be computed under IEEE double precision floats; these p-values are marked as "<1E-303" in the table.

**Supplementary Table 5.** The average, standard error of mean (SEM), and *p*-values for precision (Prec), recall (Rec) and F1-score (F1) for the agreement between manual pairwise protein alignment and automatic structure alignment by US-align, SPalign, SSM, MICAN, and Dali on the MALIDUP dataset.

| Methods | Prec mean | Prec SEM | Prec *p*-value | Rec mean | Rec SEM | Rec *p*-value | F1 mean | F1 SEM | F1 *p*-value |
|---|---|---|---|---|---|---|---|---|---|
| US-align | 0.741 | 1.48E-2 | * | 0.657 | 1.59E-2 | * | 0.782 | 1.45E-2 | * |
| SPalign | 0.720 | 1.62E-2 | 3.28E-1 | 0.634 | 1.70E-2 | 3.23E-1 | 0.757 | 1.58E-2 | 2.39E-1 |
| Dali | 0.702 | 2.06E-2 | 1.21E-1 | 0.621 | 2.03E-2 | 1.64E-1 | 0.709 | 2.05E-2 | 3.85E-3 |
| SSM | 0.601 | 2.23E-2 | 2.93E-7 | 0.519 | 2.16E-2 | 4.51E-7 | 0.615 | 2.26E-2 | 1.17E-9 |

* The *p*-values are from paired Student's t-tests against US-align.

**Supplementary Table 6.** The average, standard error of mean (SEM), and *p*-values for TM-score$_{RNA}$ (TM), RMSD (RMS), and alignment coverage (COV) in multiple RNA alignment by US-align, Matt, LinearTurboFold, and MAFFT-xinsi for all 31 groups of RNAs.

| **Methods** | **TM mean** | **TM SEM** | **TM *p*-value** | **RMS mean** | **RMS SEM** | **RMS *p*-value** | **COV mean** | **COV SEM** | **COV *p*-value** |
|---|---|---|---|---|---|---|---|---|---|
| US-align | 0.714 | 0.029 | * | 2.72 | 0.18 | * | 0.907 | 0.013 | * |
| Matt | 0.679 | 0.029 | 3.22E-4 | 3.28 | 0.34 | 6.79E-2 | 0.874 | 0.018 | 2.12E-3 |
| LinearTurboFold | 0.619 | 0.040 | 6.24E-5 | 11.54 | 1.78 | 1.38E-5 | 0.961 | 0.006 | 2.73E-6 |
| MAFFT-xinsi | 0.636 | 0.036 | 3.61E-5 | 11.31 | 1.86 | 4.50E-5 | 0.953 | 0.008 | 4.80E-6 |

* The *p*-values are from paired Student's t-tests against US-align.

**Supplementary Table 7.** The average, standard error of mean (SEM), and *p*-values for TM-score (TM), RMSD (RMS), and alignment coverage (COV) in multiple protein alignment by US-align, PROMALS3D, Matt, MAMMOTH-mult, and MUSTANG for all 92 SCOPe protein folds.

| **Methods** | **TM mean** | **TM SEM** | **TM *p*-value** | **RMS mean** | **RMS SEM** | **RMS *p*-value** | **COV mean** | **COV SEM** | **COV *p*-value** |
|---|---|---|---|---|---|---|---|---|---|
| US-align | 0.430 | 0.010 | * | 3.91 | 0.08 | * | 0.687 | 0.012 | * |
| PROMALS3D | 0.314 | 0.009 | 7.17E-15 | 9.59 | 0.36 | 6.47E-35 | 0.690 | 0.010 | 8.65E-1 |
| Matt | 0.309 | 0.012 | 1.26E-12 | 5.37 | 0.20 | 3.89E-10 | 0.469 | 0.016 | 1.92E-21 |
| MAMMOTH-mult | 0.307 | 0.011 | 2.15E-13 | 9.88 | 0.41 | 6.21E-31 | 0.680 | 0.015 | 7.01E-1 |
| MUSTANG | 0.300 | 0.012 | 2.45E-14 | 8.23 | 0.42 | 1.04E-18 | 0.596 | 0.018 | 4.68E-5 |

* The *p*-values are from paired Student's t-tests against US-align.

**Supplementary Table 8.** The average, median, standard error of mean (SEM), and *p*-values for RMSD (RMS) of RNA-protein docking by US-align, 3dRPC, and PRIME.

| **Methods** | **RMS mean** | **RMS median** | **RMS SEM** | **RMS *p*-value** |
|---|---|---|---|---|
| US-align | 63.98 | 48.93 | 2.64 | * |
| 3dRPC | 64.21 | 56.51 | 2.14 | 9.38E-1 |
| PRIME (with templates from PRIME) | 74.31 | 63.35 | 2.81 | 2.80E-8 |
| PRIME (with templates from US-align) | 71.74 | 51.28 | 2.94 | 6.86E-7 |

* The *p*-values are from paired Student's t-tests against US-align.
